# The genetic basis for synchronised time perception in plant populations

**DOI:** 10.1101/2025.02.25.640030

**Authors:** Sarah C. L. Lock, Kayla McCarthy, James Ronald, Muhammad Usman Anwer, Amanda M. Davis, Marc Hallstein, Ethan J. Redmond, Marina Knight, Seth J. Davis, Daphne Ezer

## Abstract

Synchronised developmental timing is important for ensuring crop uniformity and high yields. However, climate change is leading to crops being grown at different latitudes, varying their exposures to photoperiods over seasons and impacting developmental timings. We investigated whether the response of circadian rhythms in seedlings to changes in photoperiod would enable us to predict the timing and synchronization of flowering. Indeed, we show that the same Quantitative Trait Loci (QTLs) are associated to circadian traits in seedlings and developmental traits during bolting, using the first recombinant inbred lines (RILs) between African and European Arabidopsis lineages, spanning diverse latitudes. Two QTLs contain K-Homology Domain RNA binding proteins (KH17, KH29) and are associated with splicing variants in known flowering genes, MADS AFFECTING FLOWERING2 and 3 (MAF2, MAF3), including generating chimeric transcripts, a potential mechanism for accelerated proteome evolution. Natural variants in KH17, including in its prion-like domain, are associated with de-coupling the mean and synchronization of flowering time, enabling greater adaptation of population-level heterogeneity in developmental timings. Our results suggest that circadian traits in seedlings could be used to screen for agriculturally relevant developmental traits in mature plants, enabling efficient breeding of climate-resilient crops.

## Main

Agricultural yield is highly correlated with the timing of developmental transitions, such as bolting or flowering^1^. Effective timing of developmental transitions ensures that plant maturation occurs under optimal environmental conditions^2,3^. Yield is also correlated with how well-synchronised these transitions are within populations of plants in a field, as synchronization prevents plants from outcompeting their neighbours^4^. Uniform flowering is associated with more efficient pollination, which further increases yield^5^.

Many crops grown in temperate climates use day length (photoperiod) as an indication of seasonal change, which induces bolting and flowering^6,7^. Equatorial crops have not been selected to respond to changes in photoperiod, as there is very little seasonal variation in day length near the equator^8,9^. To mitigate the impacts of climate change on food production, farmers in temperate countries will require crops that have the climate-resilient qualities of equatorial plants, while maintaining the developmental synchronization of current temperate varieties under seasonality^10,11^. Moreover, breeding needs to occur rapidly to enable us to adapt to climate change. One strategy to accelerate breeding is to identify traits in seedlings that are strongly associated with agriculturally-relevant traits in mature plants, reducing the time and space^12,13^. We hypothesized that the ability of the seedling circadian clock to respond to the changes in day length (photoperiod) – specifically lengthening daylength that is reminiscent of the photoperiod shifts associated with spring^14^ – would correlate with flowering time, because photoperiod is a critical input into this developmental pathway^15^.

The plant circadian clock consists of multiple transcription activators and repressors that form a complex network of interlinked feedback loops, and nearly all components of the circadian clock are in some way linked to flowering. For instance, EARLY FLOWERING 3 and 4 (ELF3, ELF4) are part of the Evening Complex of the core clock and their mutants display rapid flowering under noninductive conditions^16–18^. Meanwhile GIGANTIA (GI) induces transcription of genes in the morning loop, including CIRCADIAN CLOCK ASSOCIATED 1 (CCA1) and LATE ELONGATED HYPOCOTYL (LHY), and its mutants display delayed flowering^19,20^.

We hypothesised that synchronised timekeeping on a daily scale would correlate with synchronised timekeeping at a developmental scale, which would enable us to identify selective criteria in seedlings associated with a wide range of adult developmental traits during bolting (**Fig. 1a**).We generated the first recombinant inbred lines (RILs) between African and European lineages using Tanzanian (Tnz-1; non-seasonal) and Belarusian (Ws-2; seasonal) accessions so that the RILs would display heterogenous responses to photoperiod changes (Supplementary Fig. 1 and Supplementary **Table 1**). For each of these RILs, we measured flowering time (days until bolting), as well as leaf count and biomass at bolting (Supplementary **Fig. 2-5**, Supplementary **Table 1**). Additionally, the standard error of these physiological measures was recorded, as crop uniformity is a critical agricultural trait.

**Fig. 1:**
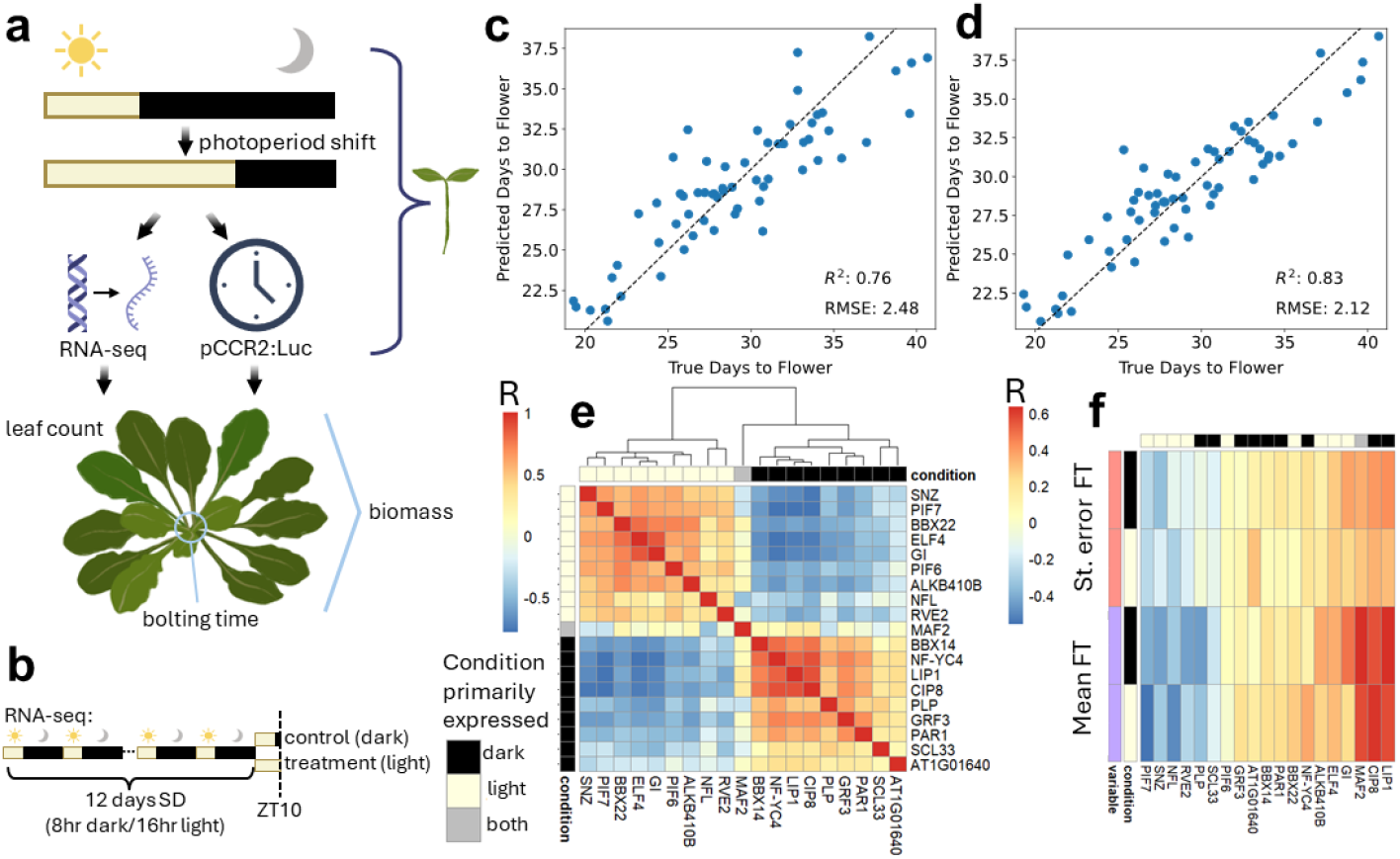
Circadian expression is predictive of flowering time. **a** This project aims to identify circadian traits in seedlings around photoperiod shifts associated with traits during bolting (which we will refer to as flowering). **b** The RNA-seq experiment performed on the RILs. The flowering time of each RIL was predicted on the basis of **c** control (dark) and **d** treatment (light) gene expression data using only genes that are differentially expressed between the two parental accessions, using leave-one-out cross validation of elastic net models. **e** Cross-correlation matrix (Pearson’s R) of the gene expression of select genes that were found as predictors in at least half of the models. Row and column annotations highlight the condition in which the genes are primarily expressed. **f** Correlation (Pearson’s R) between gene expression (from control and treatment conditions) of each of these selected genes and flowering time (FT) mean and standard error (a measure of synchronization).

We used two complementary assays to assess the response of seedlings to lengthening days (**Fig. 1a**). First, we measured gene expression at Zeitgeber Time (ZT) 10 in short day conditions (SD, 8hr light/16hr dark), either in samples collected in the dark (control) or after two-hours of extended daylight (treatment) (**Fig. 1b**, Supplementary **Table 2**). Additionally, we used this RNA-seq data to generate a high-resolution genetic map for our RILs^21^ (Supplementary **Fig. 6-8 and** Supplementary **Table 3**). As a complementary approach, we measured circadian rhythms under lengthening days. These RILs contain pCCR2:LUC, allowing us to measure luminescence over time as a marker of circadian clock activity^22^ (Supplementary **Fig. 1** and Supplementary **Table 4**). This enabled us to measure both the transcriptome-wide changes associated with increased day length at a single point in time and the response of a key circadian reporter over a time series.

Our first aim was to determine whether seedling gene expression could be used to predict adult traits, such as flowering time. After filtering for genes that have differential expression in the two parental accessions (Supplementary **Fig. 9-11**), we used an elastic net approach^23^ to select a smaller subset of genes whose expression in seedlings is predictive of adult traits, with **Fig. 1c, d** showing the predictive capacity of the model trained using either the RNA-seq data from the control or treatment conditions. Critically, the flowering time of each RIL was predicted independently using a model that was trained without that RIL (leave-one-out cross-validation) to ensure that our model was not overfitting. Similar elastic net models were developed for leaf count and biomass (Supplementary **Fig. 12, 13** and Supplementary **Table 4**), using treatment and control RNA-seq. Of the 237 genes appearing in more than half of the final models (Supplementary **Table 4**), many are known to be associated with the circadian clock, photoperiod detection, and flowering time^24^. These include key circadian genes such as *ELF4, GI* and *REVEILLE 2* (RVE2)^25^, and *PHYTOCHROME INTERACTING FACTORs 6* and *7* (PIF6, PIF7) which are associated with environment-dependent hypocotyl elongation that are regulated by light and the circadian clock^26,27^. LIGHT INSENSITIVE PERIOD1 (LIP1) is involved with photomorphogenesis in response to photoperiod^28^. COP1-INTERACTING PROTEIN 8 (CIP8) is involved in degradation of ELONGATED HYPOCOTYL5 (HY5), which is part of the light-responsive pathway^29^. Of the key genes identified, only *MADS AFFECTING FLOWERING 2* (MAF2) showed equal expression in light and dark (**Fig. 1e and** Supplementary **Fig. 14**) and its expression was only weakly correlated with the other identified genes. In contrast, *GI* and *ELF4* had highly correlated expression patterns, as do *LIP1* and *CIP8*. Expression of *LIP1, CIP8, MAF2, GI* and *ELF4* were all positively correlated with flowering time and the standard error of flowering time, while *PIF7* expression is negatively correlated with flowering time (**Fig. 1f**). These results confirm that expression in seedlings is predictive of traits at bolting. This also supports our hypothesis that the circadian clock is a pathway whose behaviour in seedlings is associated with traits at bolting.

Next, we identified genetic loci that are jointly associated with circadian response to photoperiod shifts in seedling and developmental traits during bolting. These photoperiod shifts mimic extreme versions of seasonal changes experienced in spring. For seedlings of each RIL, we measured luminescence during the following three experimental treatments (i) seven days of short day (SD, 8hr/16hr light/dark cycles), (ii) seven days of long day (LD, 16hr/8hr light/dark) (iii) 4 days of SD exposure followed by 3 days of LD exposure. We analysed these luminescence curves over time using statistical techniques from Functional Data Analysis (FDA)-- ^30^see **Fig. 2a**, Supplementary **Fig. 15 and** Supplementary **Table 4**. This enabled us to non-parametrically quantify the shapes of the curves (shape), in addition to how heterogeneous the shapes were (shape spread) – see Supplementary **Fig. 16**. We could also evaluate how much the shape changed before and after the photoperiodic shift and the heterogeneity of this shift between individual plants (shift and shift spread) – see Supplementary **Fig. 17**. Finally, we aligned the curves in the 48 hours before and after the photoperiod shift using time warping and quantified the phase changes (phase shift), which– unlike the other shift parameters– focus only on differences in phase, rather than differences in shape, before and after the photoperiod shift.

**Fig. 2:**
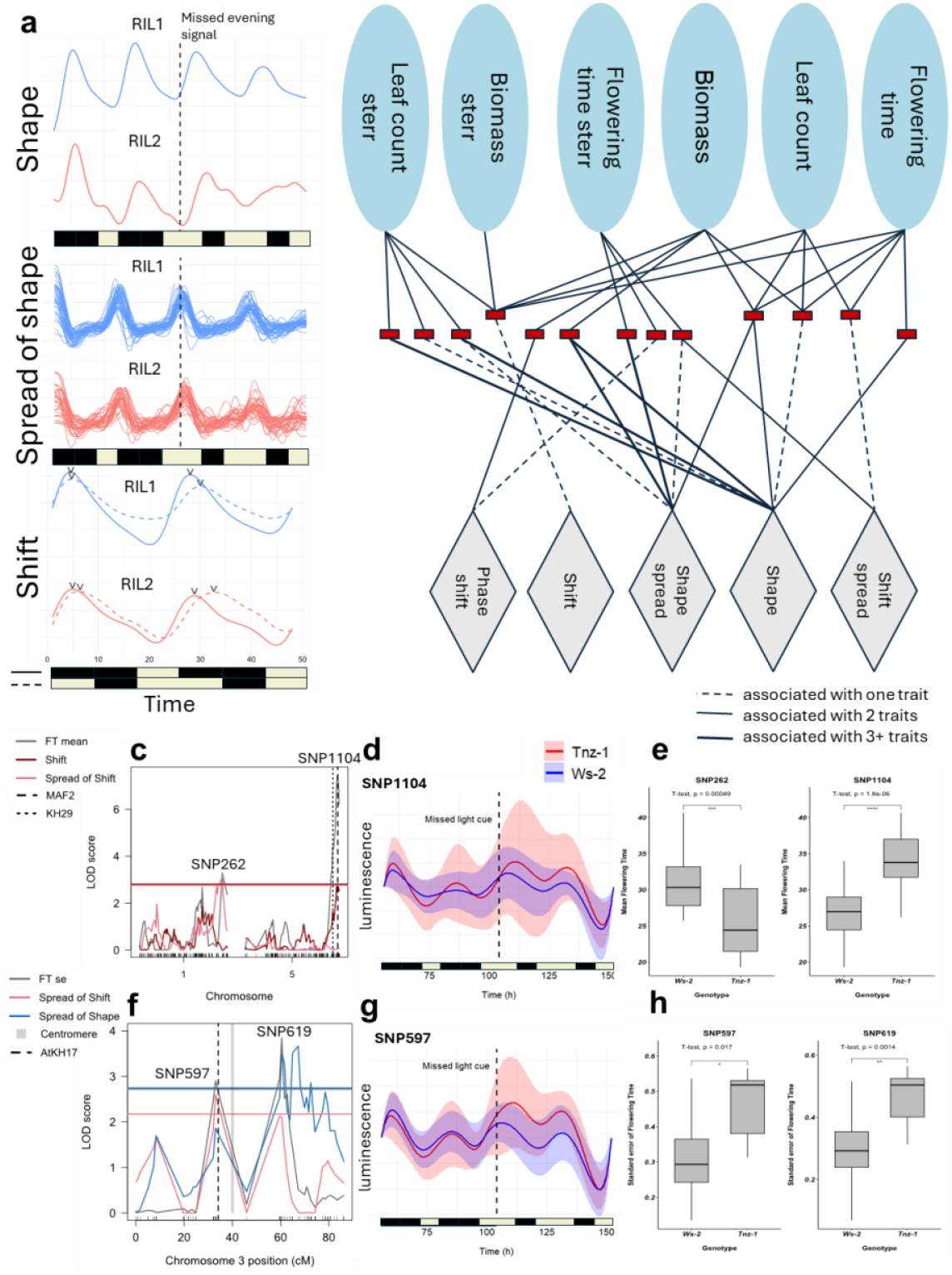
QTLs associated with response to photoperiod shifts in seedlings and bolting-associated traits. **a** Here we provide illustrative examples to explain what constitutes shape, spread and shift. For each of these parameters, we provide examples of two RILs that have divergent values of these parameters. For shape, the mean bioluminescence curve is shown for two RILs in the 48 hours before and after the photoperiod shift. For spread of shape, the individual bioluminescence curves are shown for the same time span. For shift, the mean bioluminescence curves are shown for the 48 hours immediately preceding and following the photoperiod shift, with arrows indicating peak expression. **b** Network representation of QTLs. Red rectangles represent distinct QTL blocks that link circadian traits in seedlings (grey diamonds) and bolting-associated traits (blue ovals). **c** Logarithm of odds (LOD) score profile of QTLs associated with mean flowering time. **d** Warping functions of normalised luminescence of RILs with Tnz-1 alleles and Ws-2 alleles at the SNP1104 locus during a short day to long day photoperiod shift. **e** Boxplots of mean flowering time for the two main loci for RILs containing the Ws-2 and Tnz-1 copy of the allele. **f-h** are equivalent to **c-e** but highlighting QTLs associated with the standard error of flowering time.

Many of these circadian traits were found to be correlated to the developmental traits at bolting (Supplementary **Fig. 18**). A Quantitative Trait Locus (QTL) analysis revealed 95 QTL loci shared between physiological traits in bolted plants and circadian-linked traits in seedlings (Supplementary **Table 5**). Adjacent SNP markers were condensed resulting in 13 distinct regions linking physiological and circadian traits, further supporting our initial hypothesis that circadian properties in seedlings are associated with developmental traits in adult plants (**Fig. 2b**). Given that in **Fig. 1d** genes tended to have an association both with flowering time mean and standard error, we were surprised to note that flowering time mean and standard error displayed different QTLs (Supplementary **Fig. 19)**. The main flowering time mean QTLs occurred on chromosome 1 and 5 (**Fig. 2c**), with the strongest peak containing *MAF2*—a region identified in prior QTL analyses of flowering time^31–34^. The chromosome 5 locus has a smaller secondary peak centred on a K-homologous (KH) family RNA-binding protein (KH29). The RILs were grouped by whether they had the Ws-2 or Tnz-1 allele at these loci– large effect size differences were observed between these RIL groups both in terms of bioluminescence readings after the SD to LD photoperiod shift (**Fig. 2d**) and flowering time (**Fig. 2e**). The QTLs that were primarily associated with standard error of flowering time were centred near the centromere of chromosome 3, where only 12 genes displayed non-zero expression levels (**Fig. 2f** and Supplementary **Fig. 20**). A KH family RNA-binding protein (KH17) was one of the two genes whose expression levels were associated with the presence of a Tnz-1-like allele in the region, the other gene being of unknown function (AT3G32930) (Supplementary **Fig. 20**). This QTL also had a large effect size in terms of bioluminescence (**Fig. 2g**) and flowering time standard error (**Fig. 2h**). Other QTLs were found associated with leaf count, flowering time and circadian traits (Supplementary **Fig. 21-22**) These reveal many QTLs that are associated with synchronised response to photoperiod shifts and synchronised bolting.

At least 6 of the approximately 30 KH-domain containing RNA-binding proteins in Arabidopsis have previously been found to be associated with flowering time (KH1, KH2, PEP, FLY, HEN4)^35–37^, including *FLOWERING LOCUS K* (FLK) which influences the splicing of *FLOWERING LOCUS C* (FLC)^38–41^. We hypothesized that the two KH genes within QTLs may be associated with splicing of a homologue of FLC. We found that there were two main clusters of splice variants of FLC family members (Supplementary **Table 6**). The splice junctions that have significantly different read counts between these two clusters (T-test p-value<0.05) are shown in **Fig. 3a** and are mostly found in MAF2– the FLC family member whose expression was a predictor of flowering time in **Fig. 1e**– and MAF3, a paralog that is adjacent to MAF2 along the genome. There is a dose-dependent response, with all lines that contain 3 or 4 copies of the *Tnz-1* version of KH17 and KH29 sharing the same splice pattern and almost all the lines containing four Ws-2 alleles for KH17 and KH29 having the same alternate splicing pattern (**Fig. 3a**). There are no genetic variants in MAF2 and only two variants in MAF3, neither of which flank any splice site (**Fig. 3b**). Differences in read counts mapping to each exon are visible in a genome viewer (**Fig. 3b**). Intriguingly, there are several trans-transcript splice junctions between MAF2 and MAF3 that would form chimeric transcripts. Three of these trans-gene splice junctions are supported by more than 8,000 independent reads across the RNA-seq experiments performed (Supplementary **Table 6**). Chimeric MAF2-MAF3 has been observed in previous studies^42,43^, and there have been genomic insertions of MAF3 exons within MAF2 in several Arabidopsis ecotypes^42^. These chimeric transcripts have the MADS-domain of the MAF2 gene and the K-box domain of MAF3, which is associated with dimerisation^44^. The level of chimeric transcripts is dose-dependent on the KH29 variant, but the causal link here is unclear due to genetic linkage between MAF2 and KH29 (**Fig. 3c**). Nevertheless, we find a consistent pattern that KH allele differences between Ws-2 and Tnz-1 correlate with MAF2 and MAF3 splicing patterns, including trans-gene formation, which is then correlated with flowering time, in a pathway that is independent of ELF4 and GI (**Fig. 3d**). While KH29 and MAF2 are found in QTLs associated with flowering time, KH17 was associated with flowering time standard error. We found that the same genes whose expressions were positively correlated with flowering time were also correlated with standard error and vice versa (**Fig. 1f and** Supplementary **Fig. 23**). If KH17 specifically impacts flowering time standard error, we hypothesized that genetic variability in this gene would de-couple the positive association between flowering time mean and standard error. To test this hypothesis, we analysed KH17 transcript sequences from the 1001 genome project (Supplementary **Table 7**)^45^. KH17 has an unstructured prion-like domain (similar to atrophin-1), which is the primary region containing variants across the ecotypes (**Fig. 4a, b**). KH29 has fewer variants across the ecotypes, but these are also primarily located in a disordered region (**Fig. 4c, d**). After clustering the ecotypes by the variant combinations (**Fig. 4e**), we mapped the distribution of variant combinations over Eurasia and North Africa (**Fig. 4f** and Supplementary **Table 7**), demonstrating a longitudinal pattern rather than a latitudinal pattern. We selected six ecotypes that had very similar KH29 sequences but different KH17 sequences. We find that although all three ecotypes with the Ws-2-like KH17 sequences have earlier flowering time, they all have high standard errors in flowering time (**Fig. 4g, h**), confirming our hypothesis that KH17 alleles may be associated with a decoupling of flowering time mean and standard error.

**Fig. 3:**
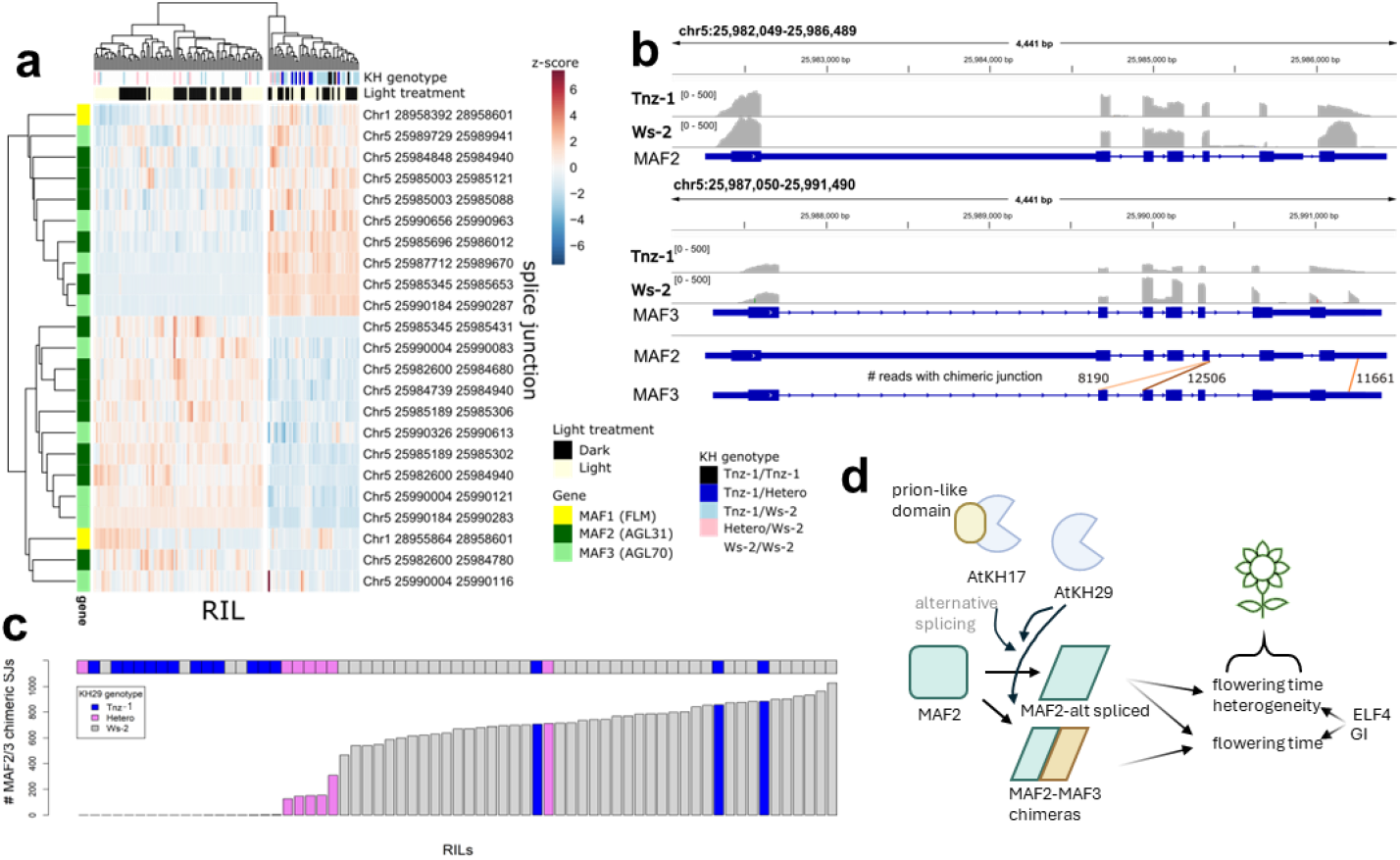
KH alleles associated with slice variants in MAF2 and MAF3. **a** There are two clusters of RNA-seq samples that have different proportions of splice junctions within FLC-family genes. Only splice junctions that account for >0.1% of total splice junctions found and that are differentially found in the two clusters (t-test, p-value<0.05) are shown. **b** Visualisation of mapped Ws-2 and Tnz-1 reads over MAF2 and MAF3. Chimeric splice junctions that are supported by more than 8,000 reads across all RILs. **c** Plot of read counts that are chimeric between MAF2 and MAF3 for each RNA-seq sample, with the horizontal bar colour-coded by KH29 genotype. **d** Proposed mechanism by which KH-proteins, ELF3 and ELF4 may influence flowering time mean and standard error.

**Fig. 4:**
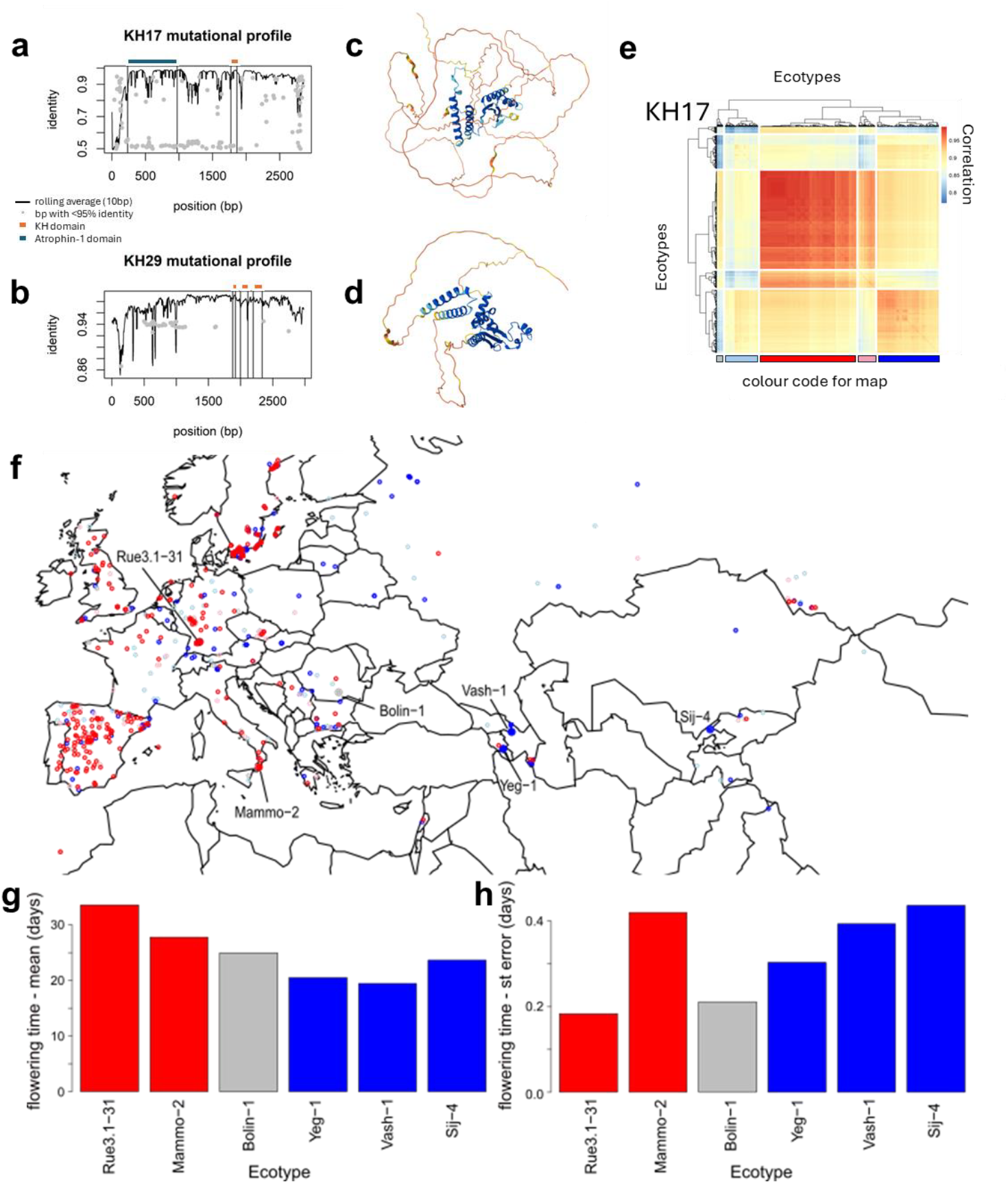
Natural variation in KH17 associated with decoupling mean flowering time and degree of synchronization. Based on a sequence alignment of all ecotypes in the 1001 genome project, we calculated the mutational profile of KH17 (**a**) and KH29 (**b**) showing the rolling average (over 10bp windows) of the percent identity–including Ns and gaps– and dots highlighting mutational hotspots, bases with less than 95% identity, ignoring Ns and gaps. Conserved domains are annotated. **c-d** Alphafold reveals low confidence structure predictions outside the core KH domain. **e** A clustering of variant co-occurrence across natural ecotypes, with clusters colour-coded, to be used as a colour-key in the map in **f**, with ecotypes whose flowering times have been measured shown with larger labelled dots. Their mean flowering times **g** and standard error of flowering times **h** are shown, with bars colour-coded by the version of KH17. Note that blue corresponds to Ws-2-like variants.

The circadian clock has been linked with lifespan in plants, animal models and humans^46,47^. Our study provides a unified mechanism of plant timekeeping across wide temporal scales. KH genes may be associated with splice variants in MAF2 and MAF3, including formation of chimeric transcripts, similar to how FLK (another KH protein) contributes to alternative splicing of FLC (another MAF family member)^36,48^. KH17 may be involved in de-coupling regulation of flowering time mean and standard error, possibly through variants in an unstructured prion-like domain, which have recently been implicated with environmental responsiveness^49^. Variation in prion-like domains and formation of chimeric genes are two strategies for rapid proteome evolution, enabling the adaptation of life cycles to diverse environmental conditions^50^. Moreover, our work suggests that photoperiod-associated transcriptional changes in seedlings are correlated with developmental traits during bolting, suggesting a strategy by which lines may be screened at the seedling stage to accelerate breeding of key traits of mature plants.

## Acknowledgements

We thank Jason Daff, Paul Scott, Harry Stevens and the rest of the University of York Horticulture team. The Viking Cluster was used in this project, which is a high-performance compute facility provided by the University of York. The authors are grateful for computational support from the University of York High Performance Computing service, Viking and the Research Computing team. Also, we thank Professor Kanchon Dasmahapatra ideas on narrative structure. Alongside Katie Kindleysides for her graphic design of Arabidopsis images.

This work was supported by funding from the Biotechnology and Biological Sciences Research Council (BBSRC)—DE, SJD, MK: BB/V006665/1. The authors also acknowledge BBSRC White Rose DTP studentships (BB/M011151/1 and BB/T007222/1) to JR (Ref.: 1792522) and EJR (Ref.: 2444228), and Royal society funding to SJD (IF\R2\2320049).

## Author contributions

S.C.L.L.: conceptualisation, methodology, data collection, software, analysis, investigation, writing – original draft, and visualization. K.M: data collection. J.R.: writing – review and editing. M.U.A: methodology, material development, writing – review and editing. A.M.D: methodology, material development. M.H: methodology, material development. E.J.R: analysis, data collection. M.K: conceptualization, methodology, writing – review and editing, supervision, and funding acquisition. S.J.D.: conceptualization, writing – review and editing, supervision, and funding acquisition. D.E.: conceptualization, methodology, analysis, writing – original draft, writing – review and editing, supervision, and funding acquisition.

## Additional information

### Supplementary Information

See Supplementary document

### Data availability

Raw and processed sequencing data have been deposited in the NCBI Gene Expression Omnibus database under accession number GSE286355. Scripts used to produce the figures have been deposited in a GitHub repository (https://github.com/scllock/SyncTimePerceptionPlants).

