## Supplementary Figures for "The genetic basis for synchronised time perception in plant populations"

Supplemental Tables:

Table S1: Information about the parental lines of the RILs and physiological data for the RILs

Table S2: Gene expression data for RILs under control (ZT10, dark) and treatment (ZT10, light)

Table S3: genetic map for RILs, along additional data about the quality of the genetic map and gene list per SNP and the VCF files used to generate the map

Table S4: shape, shape spread, shift, shift spread, and phase shift parameters extracted for each RIL, alongside the parameters used in the elastic nets for each of the physiological parameters, the resulting coefficients for each elastic net and a list of the genes that were found in multiple elastic net models.

Table S5: Summaries of all QTL analysis for every circadian and physiological parameter tested.

Table S6: Splice variants for all RILs, with trans-splice variants in MAF2-MAF3 specifically indicated.

Table S7: Sequences of KH17 and KH29 genes from 1001 genome project, alongside data of location for each ecotype and which cluster it came from.

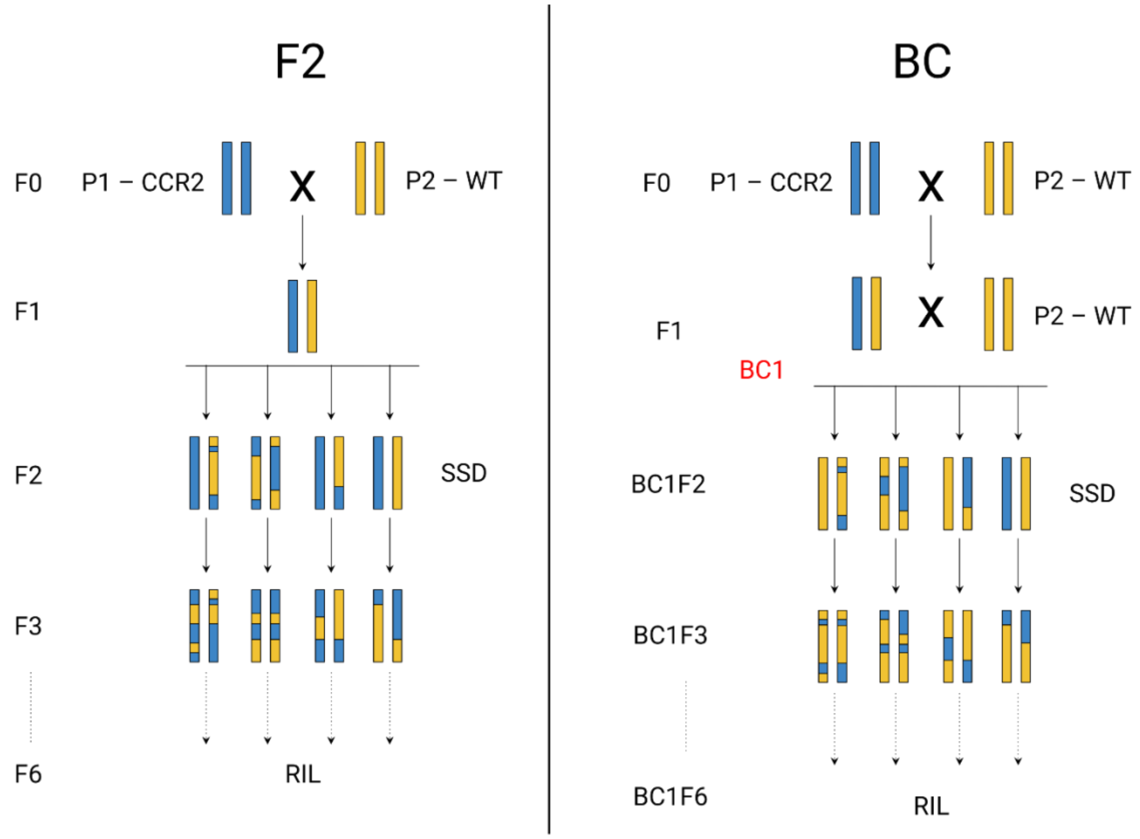


**Supplementary Fig. 1**: Schematic diagram representing the crossing strategies used to generate the recombinant inbred line (RIL) population. Within the RIL population approximately 58% were made using the F2 strategy and 42% using back-crossed (BC). For the F2 sub population Tnz-1 is crossed with Ws-2 to produce heterozygous F1 individuals. F1 individuals are then selfed to produce the F2 population which has genetic recombination between the two parent genomes. The F2 individuals then undergo single seed decent (SSD) for 6 generations leading to homozygous individuals. For the BC sub population Tnz-1 is crossed with Ws-2 to produce heterozygous F1 individuals. Then the F1 individuals are crossed back to the Ws-2 parent, this results in a back cross (BC) population BC1. The BC1 individuals then undergo 6 generations of SSD to attain homozygous individuals.

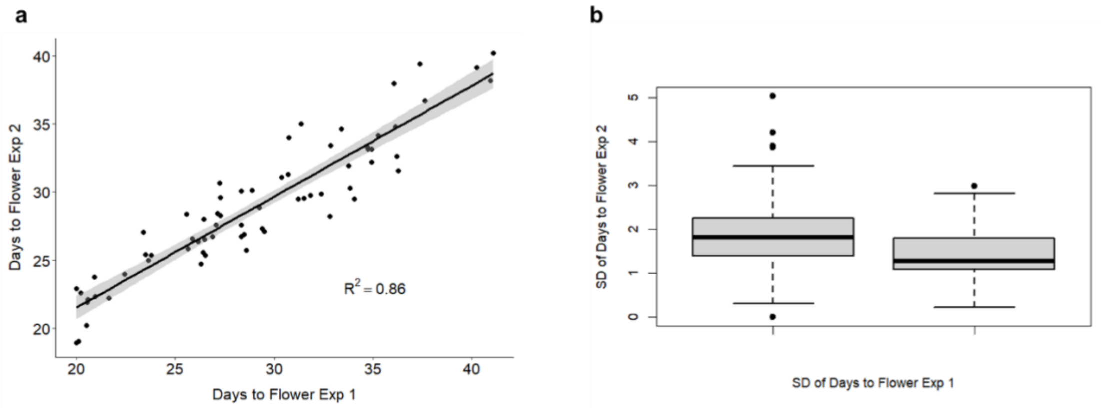


**Supplementary Fig. 2**: Relationship and variability in flowering time across two replicate experiments. **a** Scatterplot showing the correlation between days to flower in Experiment 1 and Experiment 2, with a fitted linear regression line (black) and 95% confidence interval (grey shading). The $R^2$ coefficient of 0.86 indicates a strong positive correlation between the two experiments. The linear regression model (mean days to flower ~ standard error) is significant ($F(1, 65) = 410$, $p = 2.2e-16$). **b** Boxplot comparing the standard deviation (SD) of days to flower between Experiment 1 and Experiment 2. The median SD values and interquartile ranges (IQRs) are displayed, with outliers represented as points beyond the whiskers.

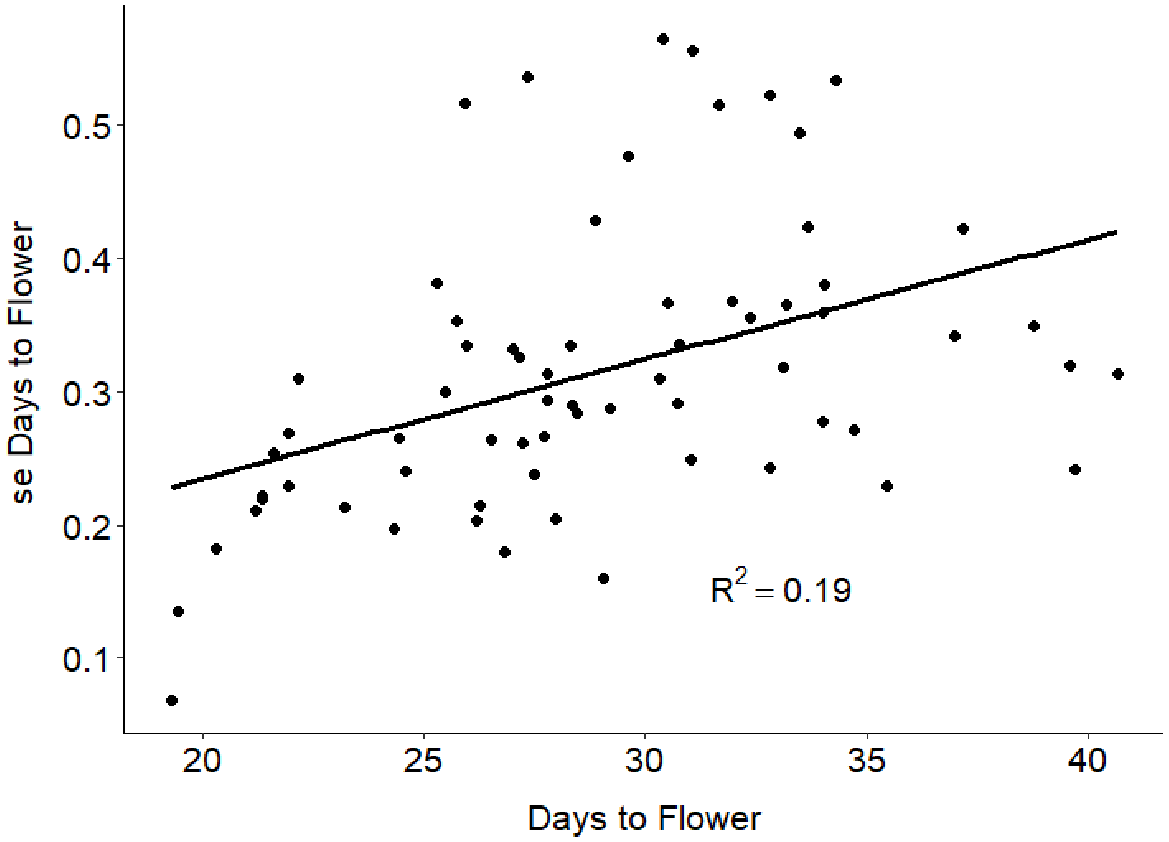


**Supplementary Fig. 3**: Scatterplot showing the correlation between between mean days to flower of combined experiments and standard error of flowering time for combined experiments, with a fitted linear regression line (black). The $R^2$ coefficient of 0.19 indicates weak correlation between the two variables. The linear regression model (mean days to flower ~ standard error) is significant ($F(1, 65) = 14.76$, $p = 0.00028$).

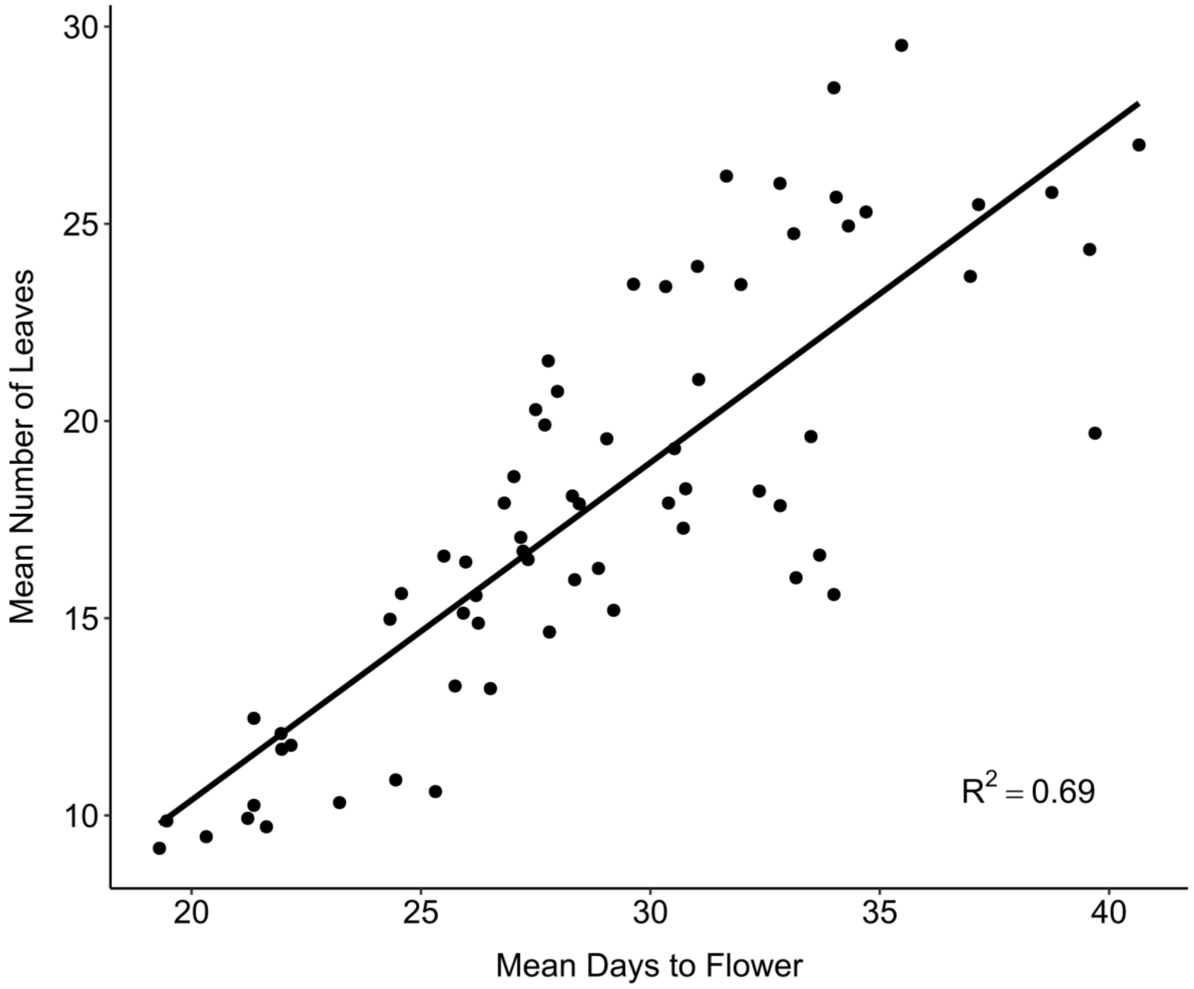


**Supplementary Fig. 4**: Scatterplot showing the correlation between between mean number of leaves of combined experiments and mean days to flower for combined experiments, with a fitted linear regression line (black). The $R^2$ coefficient of 0.69 indicates strong correlation between the two variables. The linear regression model (mean leaves ~ mean days to flower) is significant (F(1, 65) = 142.3, p = 2.2e-16).

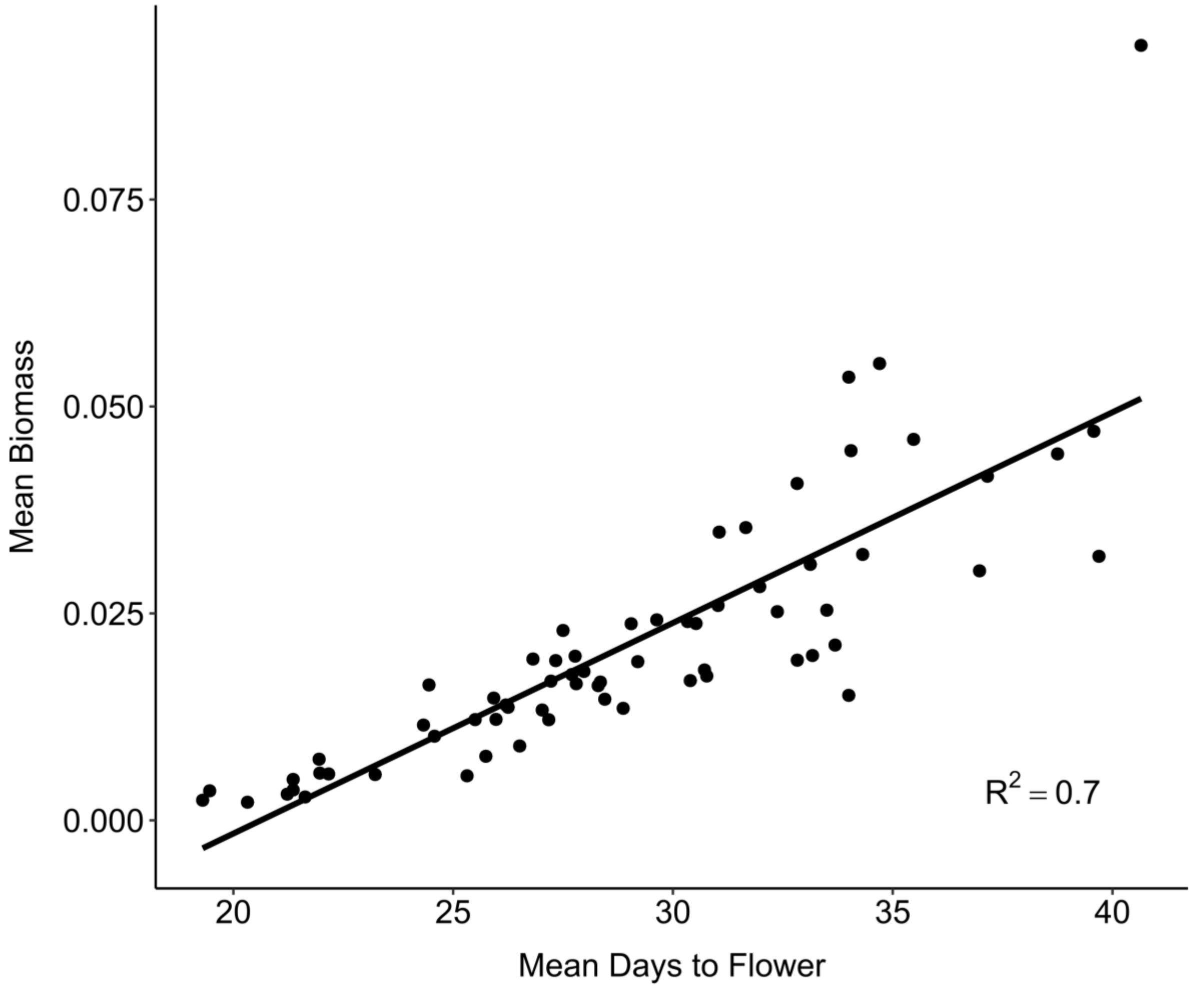


**Supplementary Fig. 5**: Scatterplot showing the correlation between between mean biomass of combined experiments and mean days to flower for combined experiments, with a fitted linear regression line (black). The $R^2$ coefficient of 0.7 indicates strong correlation between the two variables. The linear regression model (mean biomass ~ mean days to flower) is significant ($F(1, 65) = 149.9$, $p = 2.2e\text{-}16$).

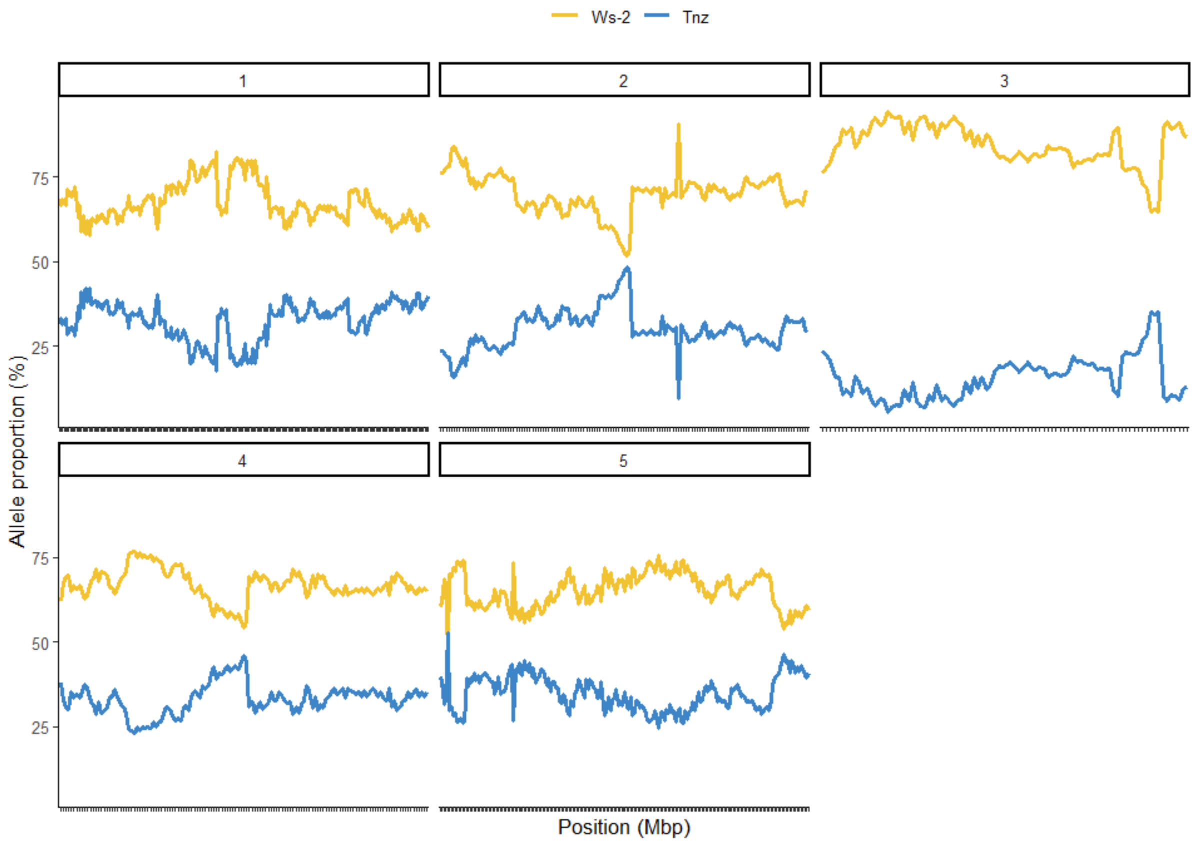


**Supplementary Fig. 6:** Allele distribution for the 789 markers along the five chromosomes. Orange and blue colours indicated the Ws-2 and the Tnz-1 allele percentages, respectively.

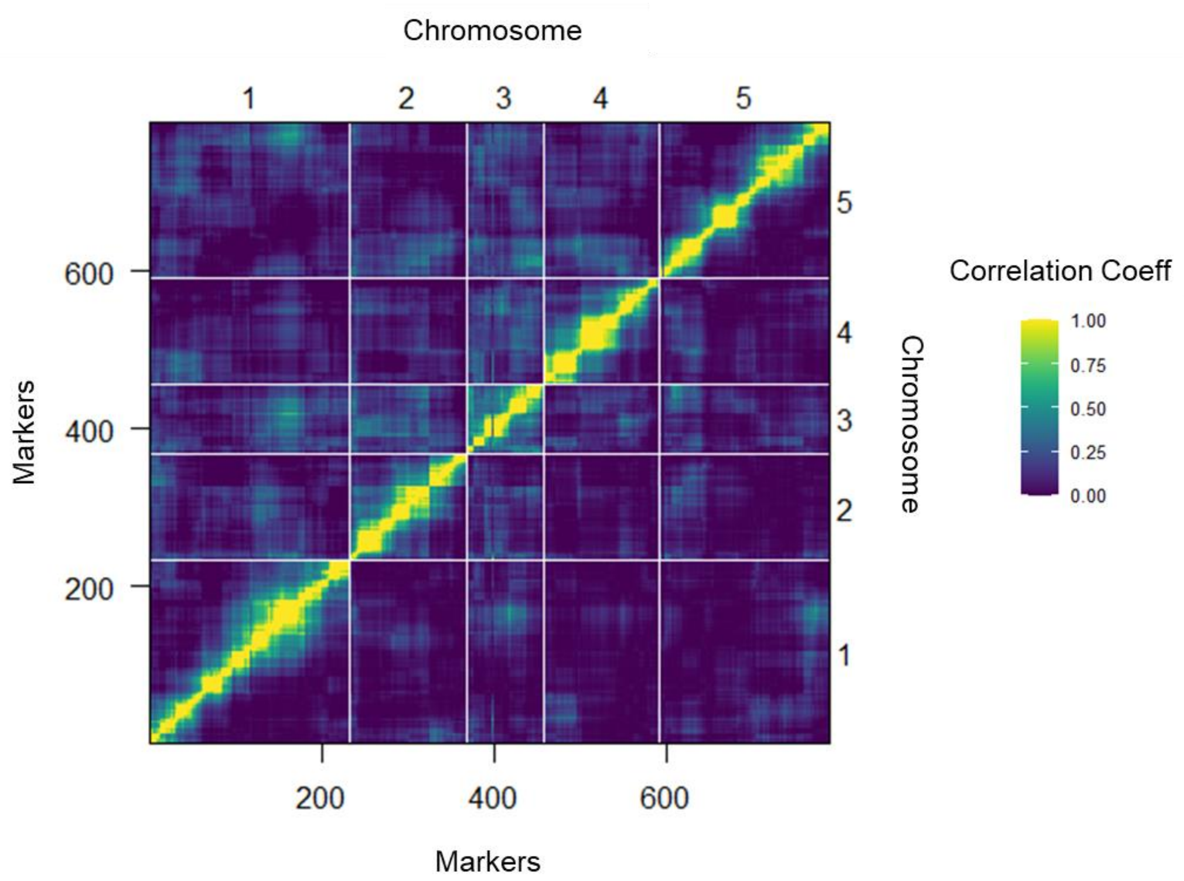


**Supplementary Fig. 7**: Pairwise marker linkage analysis. The estimated recombination fraction and Logarithm of the Odds (LOD) scores for all pairs of markers are shown in the upper-left and lower-right triangle, respectively. High correlation between markers indicates marker linkage (yellow) while the blue colour shows low correlation values indicating unlinked markers. The grid delineates the five chromosomes.

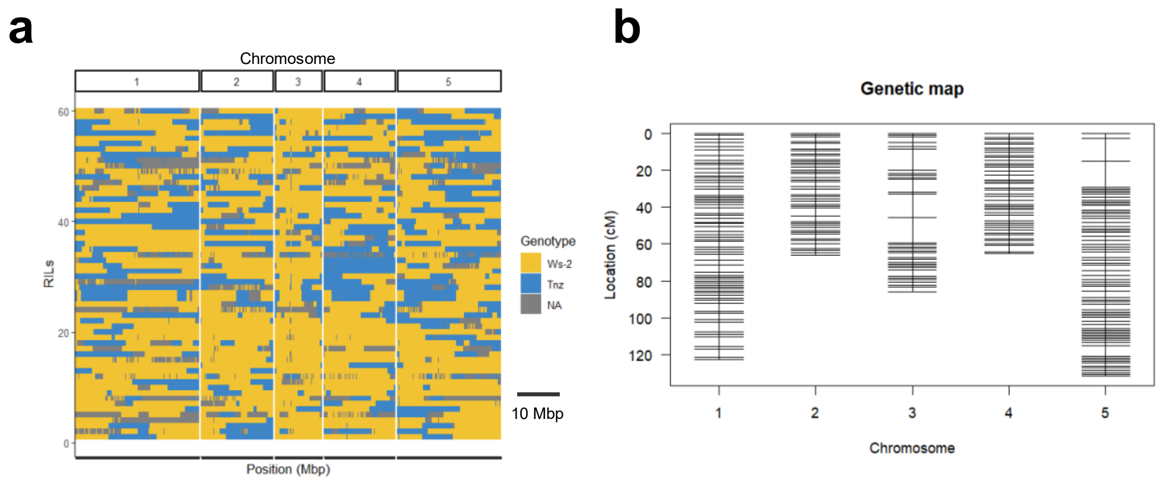


**Supplementary Fig. 8: a** Haplotype representation of the 63 RILs. Each row corresponds to a RIL. Columns represent the 789 genetic markers physically anchored on the five chromosomes. Orange boxes indicate Ws-2 genotype, and blue boxes indicate Tnz-1 genotypes. Grey boxes show the missing data. **b** Genetic linkage map of *A. thaliana*. Black bars indicate SNP markers.

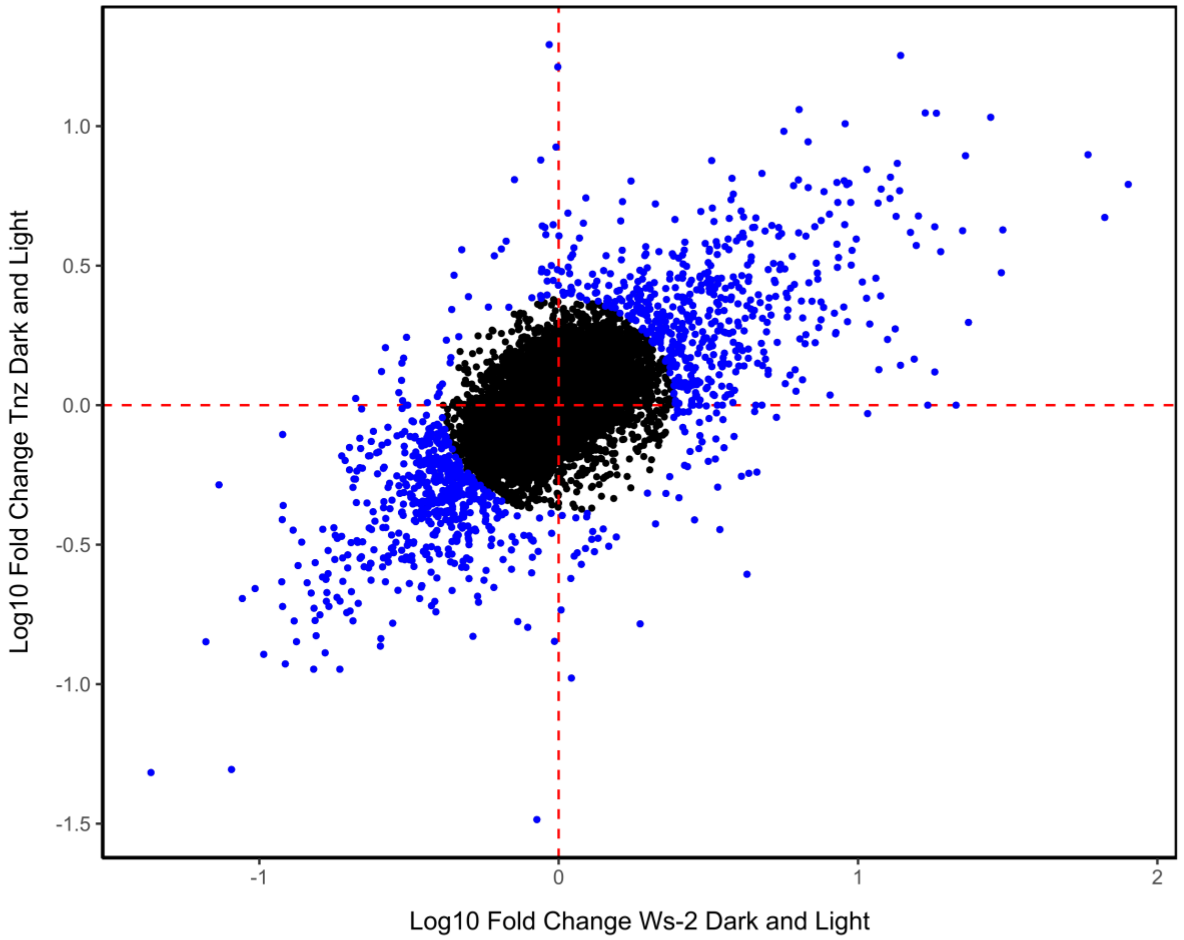


**Supplementary Fig. 9**: Log fold change between Ws-2 under dark and light conditions vs log fold change between Tnz-1 under dark and light conditions from RNA-seq experiment using whole plant tissue. TPM values were filtered to be >10. Red dashed lines show X = 0 and Y = 0. Blue points indicate genes that are in the top 20% of Euclidean distances from the centre. Genes in blue were used in Elastic net analysis.

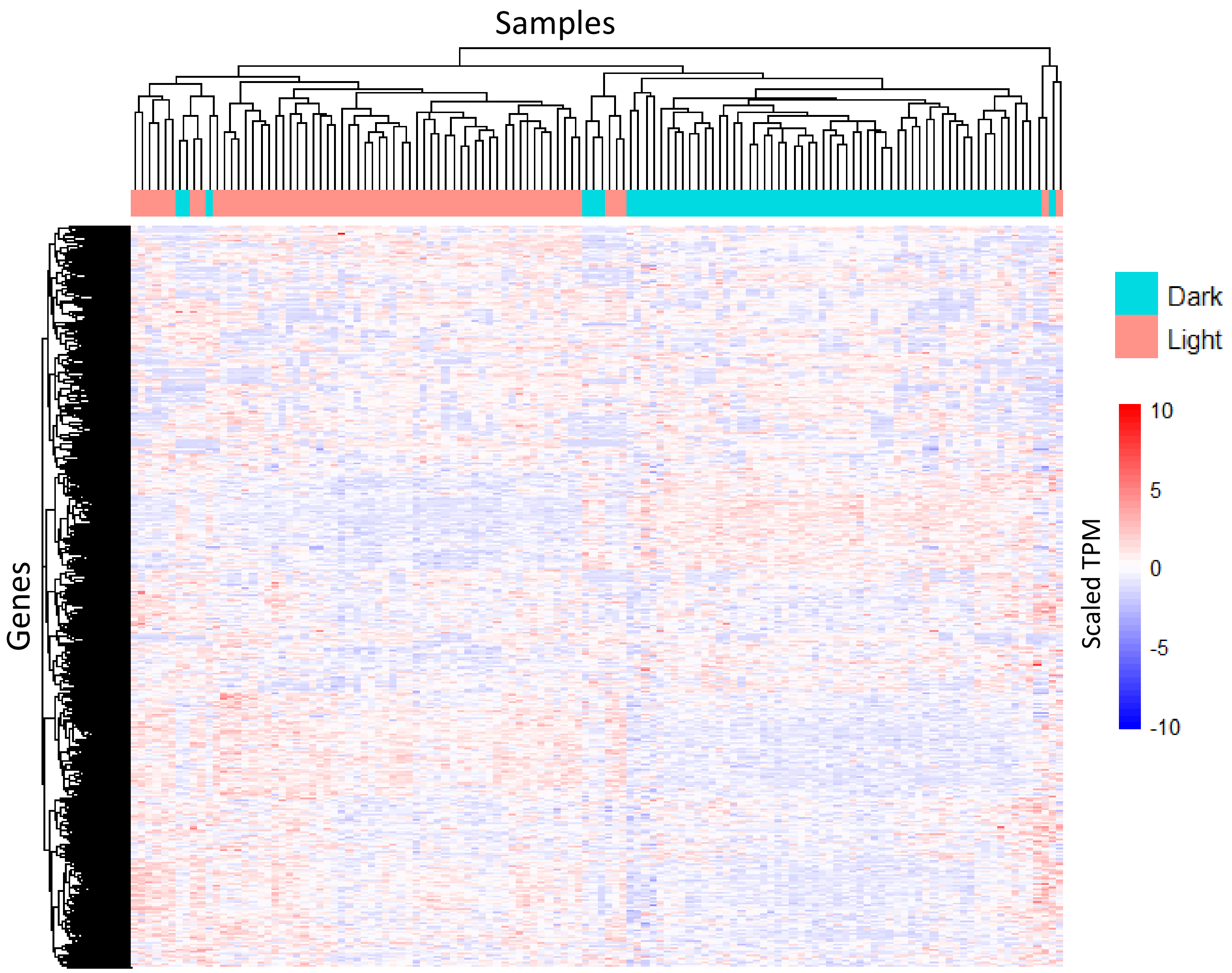


**Supplementary Fig. 10:** Gene expression values across all samples for genes where 50% of samples contained TPM values > 10. Hierarchical clustering across each sample shows that plants with the same condition status often cluster together. For all hierarchical clustering, we used the ‘complete’ linkage method.

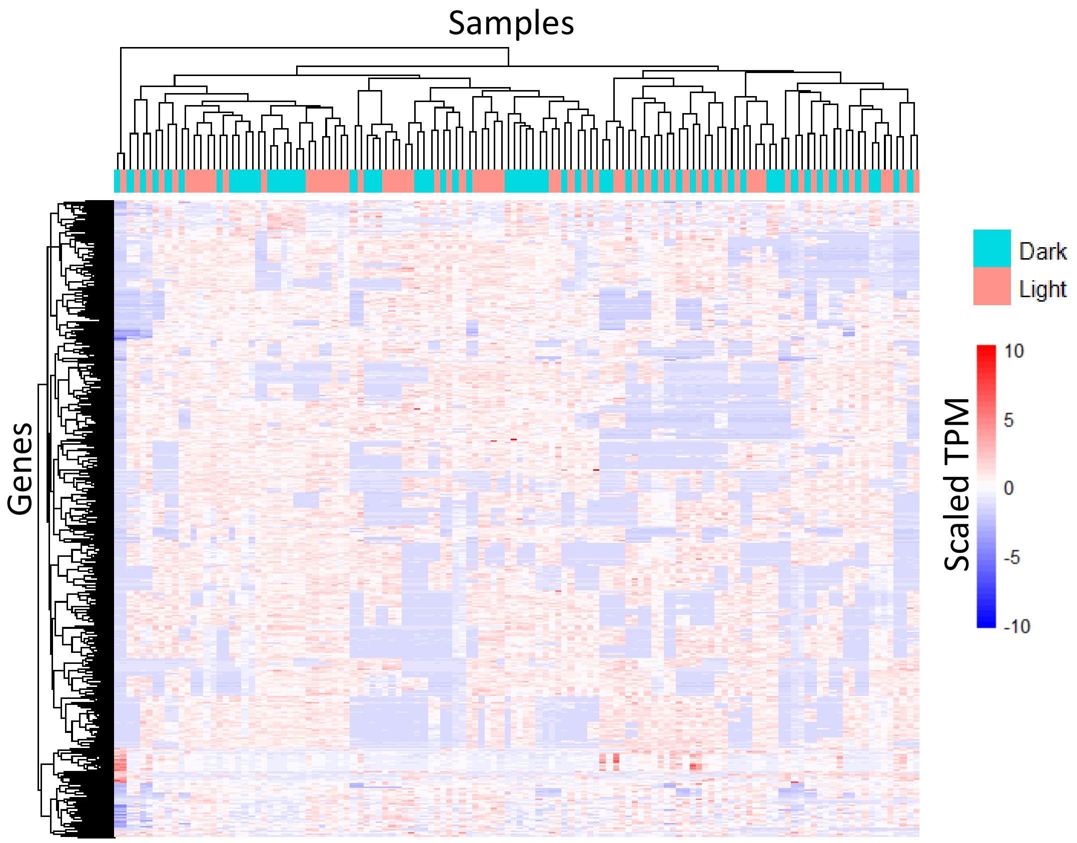


**Supplementary Fig. 11**: Gene expression values across all samples for genes where 50% of samples contained TPM values > 10. Only genes which were identified as having a Euclidean distance in the top 10% when comparing log fold change of ecotype (blue points in Supplementary **Fig. 9**) are shown (log fold change Ws-2 in Dark and Light vs log fold change Tnz-1 in Dark and Light). Hierarchical clustering across each sample shows that plants with the same condition status no longer cluster together.

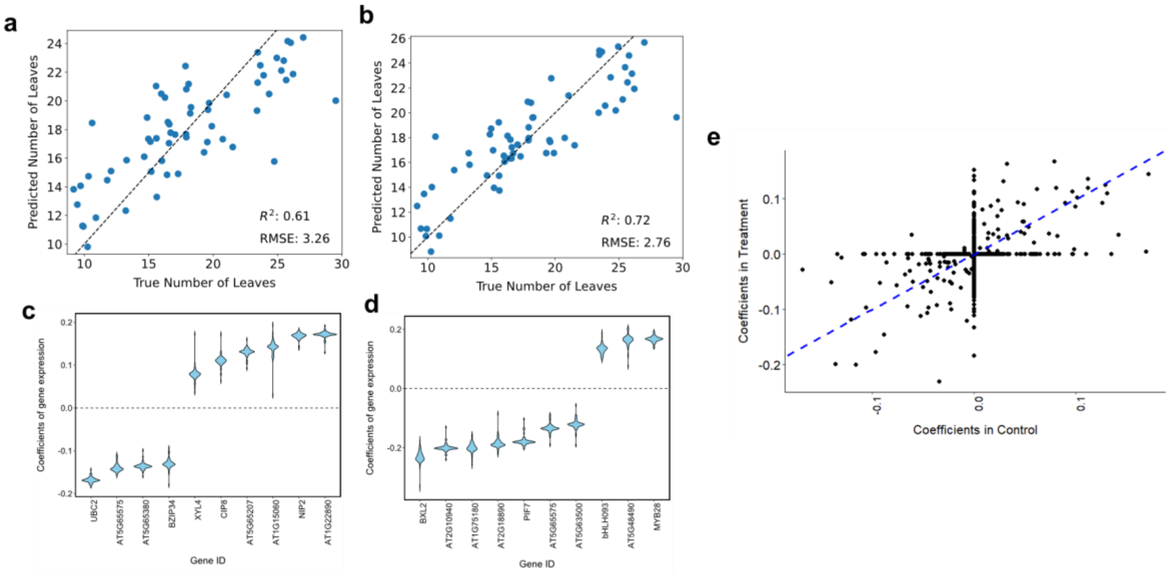


**Supplementary Fig. 12**: The number of leaves at the time of flowering of each RIL was predicted based on control (dark) (**a**) and treatment (light) (**b**) gene expression data using only genes that are differentially expressed between the two parental accessions, using leave-one-out cross validation of elastic net models. The coefficients from elastic net models for the top 10 highest absolute coefficients for control (dark) (**c**) and treatment (light) (**d**). **e** Scatter plot showing coefficients from the elastic net where 50% of models contained non-zero coefficients in both treatment (dark) vs control (light). The blue dashed line is y=x.

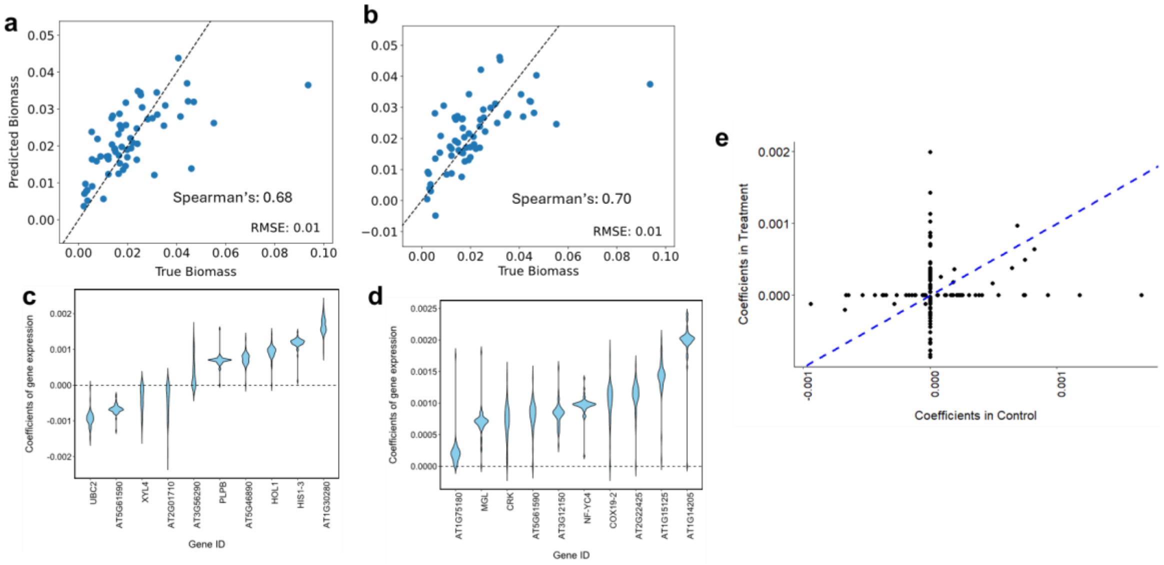


**Supplementary Fig. 13**: The dry biomass at the time of flowering of each RIL was predicted on the basis of control (dark) (**a**) and treatment (light) (**b**) gene expression data using only genes that are differentially expressed between the two parental accessions, using leave-one-out cross validation of elastic net models. The coefficients from elastic net models for the top 10 highest absolute coefficients for control (dark) (**c**) and treatment (light) (**d**). **e** Scatter plot showing coefficients from the elastic net where 50% of models contained non-zero coefficients in both treatment (dark) vs control (light). The blue dashed line is y=x.

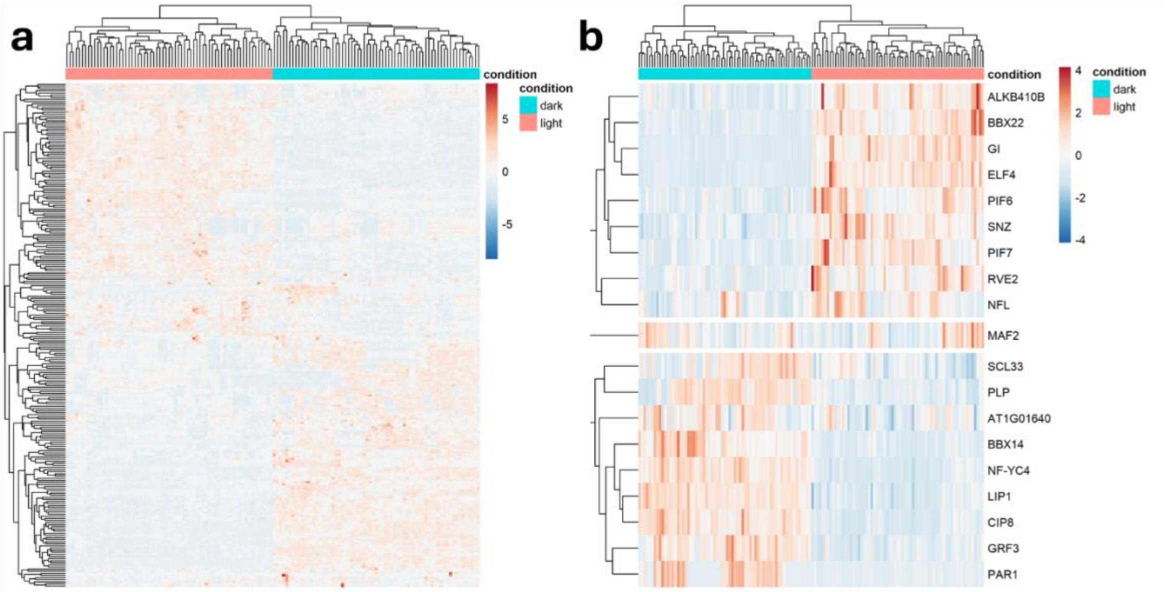


**Supplementary Fig. 14: a** A heatmap shows the gene expression levels (z-score of TPMs) of all genes that were found in at least 3 of the 6 Elastic Net models constructed. **b** A heatmap showing a selection of genes from (**a**), selected due to the importance of the genes in flowering time, the circadian clock and photoperiod detection, again showing the z-score of TPMs.

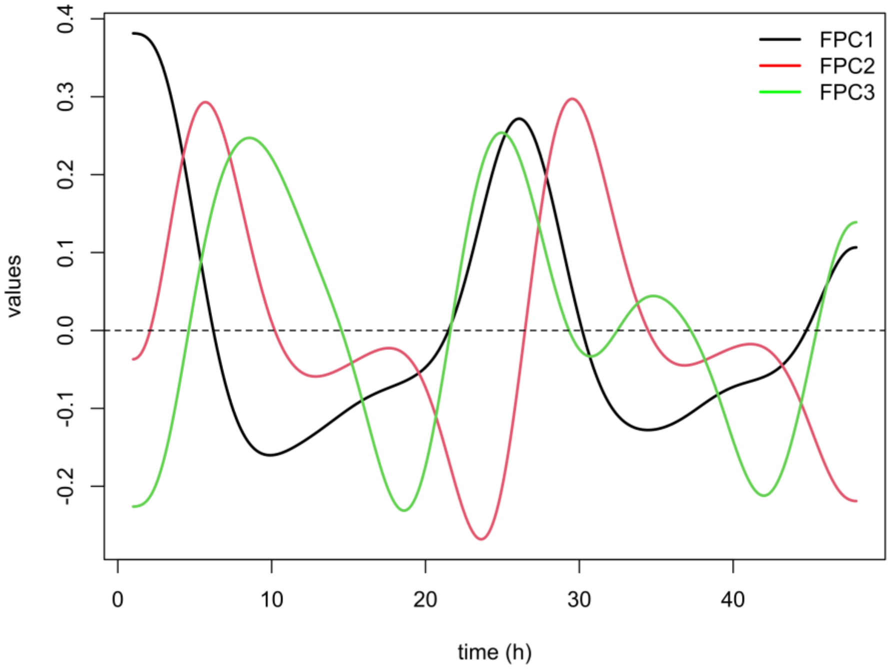


**Supplementary Fig. 15**: The first three functional Principal Components (FPCs) of the first derivative (velocity) of bioluminescence over time, in the 24 hours immediately before and after the photoperiod shift. These demonstrate that the FPCs capture information about the rate of change in bioluminescence at different times of day.

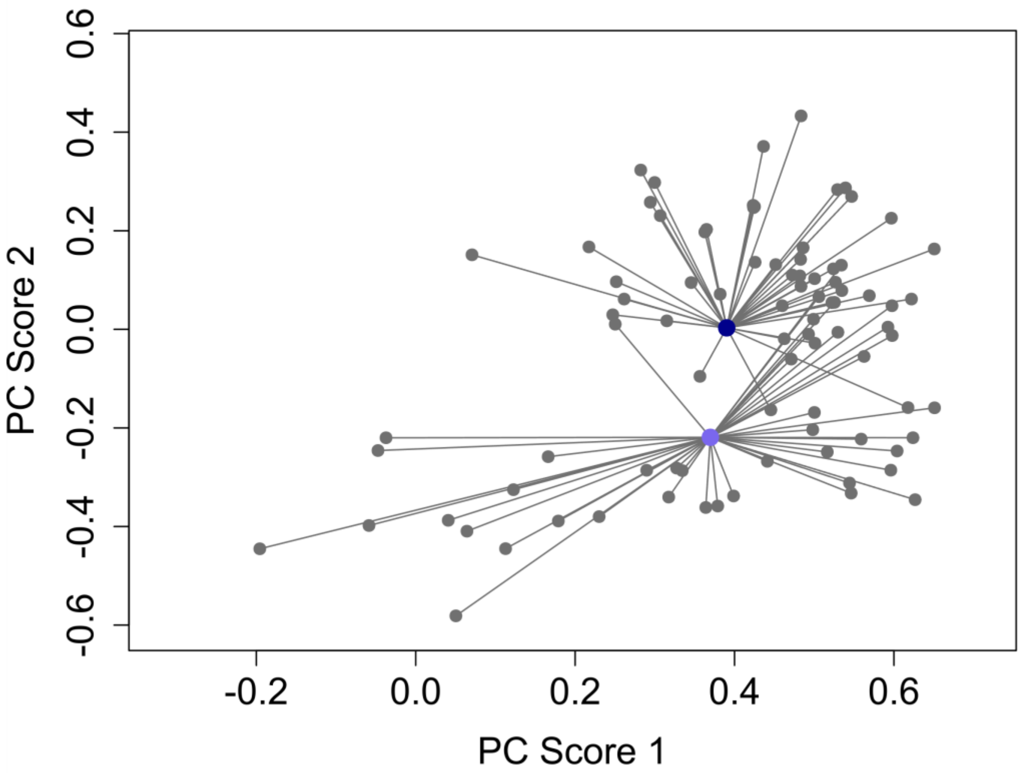


**Supplementary Fig. 16**: Here we illustrate how the **shape** and the **spread of shape** are calculated. For illustrative purposes, we are only including two randomly selected genotypes. Each grey point in this PCA plot represents a bioluminescence curve from one individual plant. The light and dark blue coloured dots are the centroids of each genotype. The PC1 and PC2 scores of the centroids are considered 'shape' parameters in our QTL analysis. The spread is the average Euclidean distance between the grey points and the centroid.

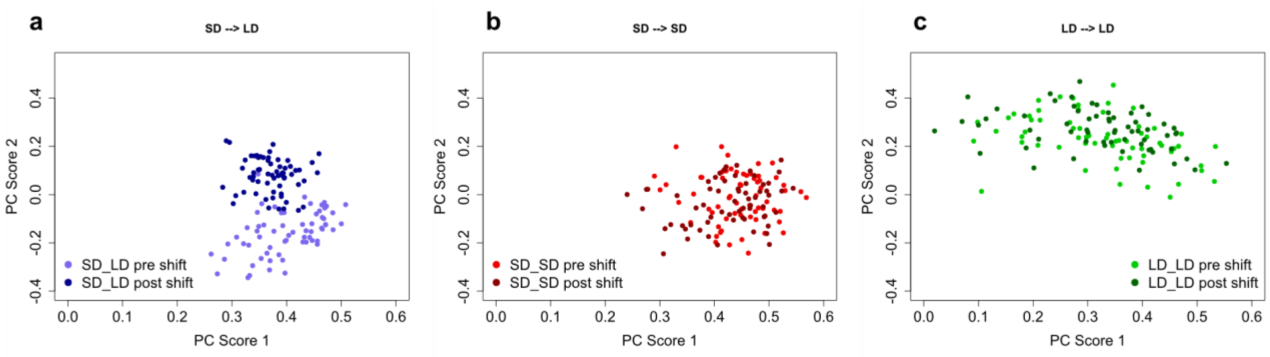


**Supplementary Fig. 17**: Functional principal component analysis (FPCA) of plants circadian responses under different day-length conditions. Each plot represents the first two principal components (PC Score 1 and PC Score 2) derived from functional data. For visual clarity, we only show the centroids—i.e. for each colour there is one point per RIL.

**a** Transition from short day (SD) to long day (LD) conditions (SD_LD). Light blue dots represent plants before the shift (48 hours pre-shift), and dark blue dots represent plants after the shift (48 hrs post-shift). **b** Short day control group (SD_SD), where plants remained under SD conditions. Light red dots represent 48 hours before the shift would have taken place, and dark red dots represent the plants in the 48 hours after the shift would have taken place. **c** Long day control group (LD_LD), where plants remained under LD conditions. Light green dots represent “pre-shift” plants, and dark green dots represent “post-shift” plants.

In **a**, we observe that there is a shift in the shape of the bioluminescence curves after the photoperiod shift, which is not present in the two controls **b** and **c** where plants were exposed to the same photoperiod conditions throughout the experiment.

Please note that the Euclidean distance between the centroids for each RIL in the 48 hours before and after the photoperiod shift is what we refer to as the ‘**shift**’ parameter in our QTL analysis. The standard error of the shifts across all individual plants is the ‘**spread in shifts’** parameter.

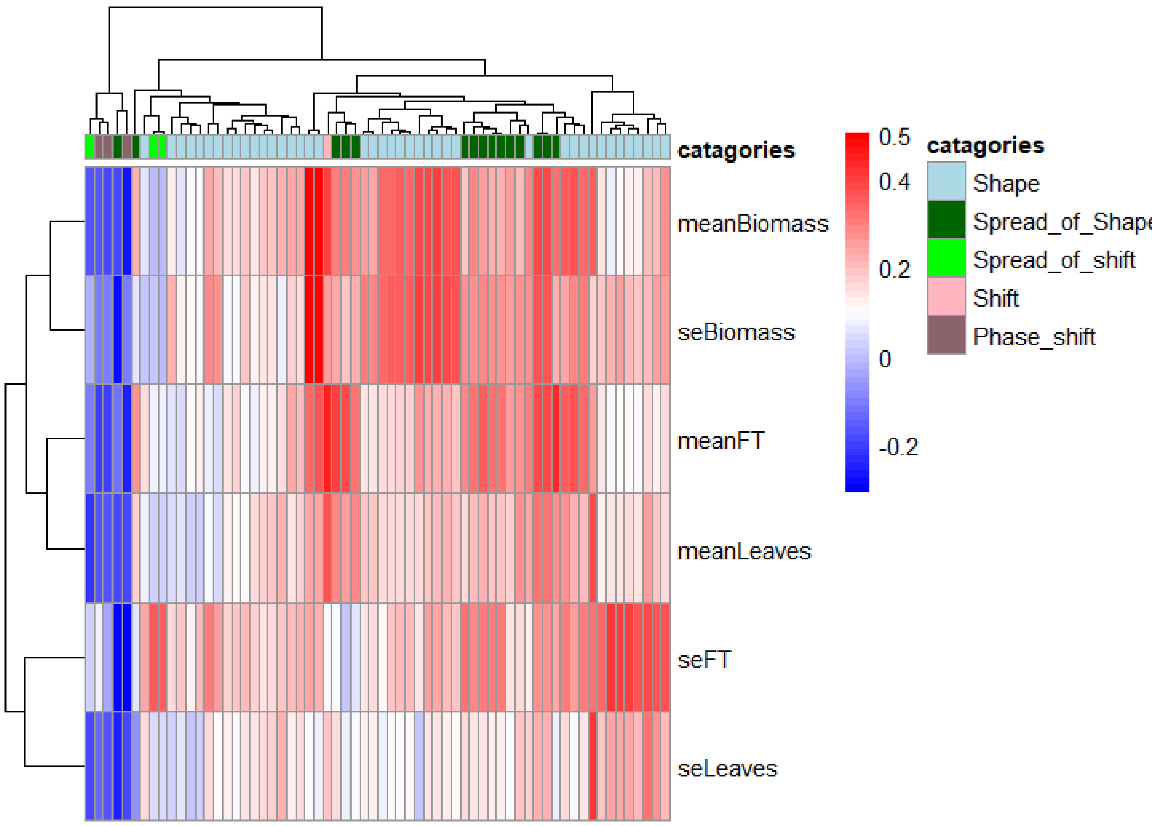


**Supplementary Fig. 18:** Heatmap showing Spearman correlations between functional traits derived from functional principal component analysis (FPCA) and developmental traits at flowering. Functional traits are categorized into five groups: Shape, Spread of Shape, Spread of Shift, Shift, and Phase Shift, indicated by color-coded annotations at the top of the heatmap. Correlation values range from -0.2 (blue, negative correlation) to 0.5 (red, positive correlation), as shown in the colour scale. The hierarchical clustering of traits and functional categories highlights relationships between specific functional features and developmental outcomes, emphasising the relationship between diurnal bioluminescence-derived metrics and developmental traits.

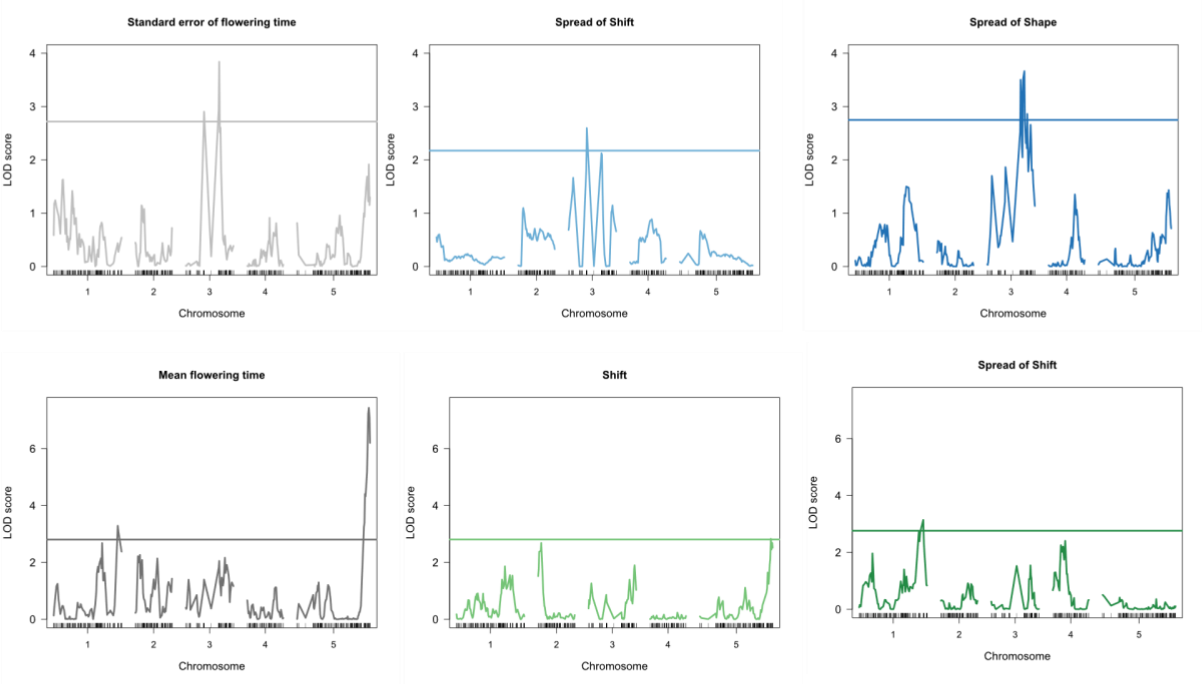


**Supplementary Fig. 19**: LOD score profile for traits depicted in **Fig. 2c,f** across all chromosomes. Horizontal lines depict the LOD score 5% threshold.

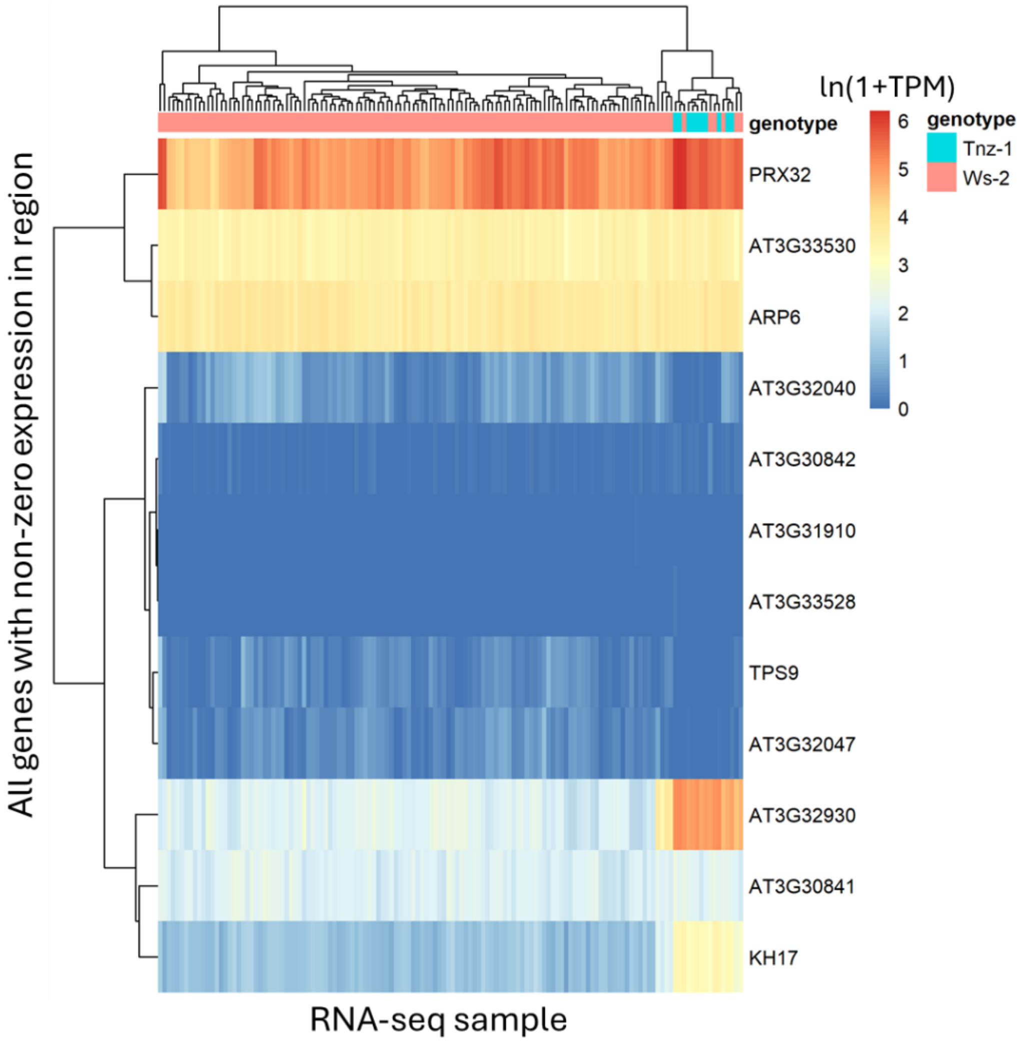


**Supplementary Fig. 20**: All genes that were in the genetic map SNP597 and the two adjacent genetic map regions were analysed. There were 63 genes in this region. Of those, only 12 of the genes had non-zero expression. We clustered the ln(1+TPM) values for these genes, annotating the allele associated with each RIL for each column. We find that KH17 and AT3G32930 are the two genes with the most expression difference between RILs with Tnz-1 and Ws-2 alleles. AT3G32930 is a protein of unknown function.

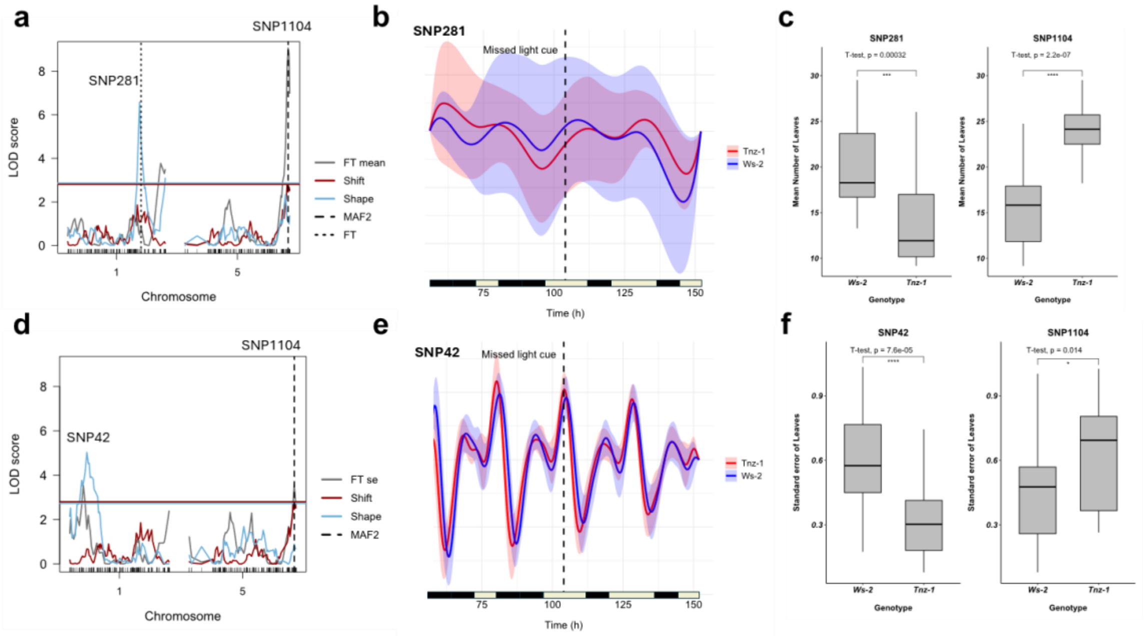


**Supplementary Fig. 21: a** LOD score profile of QTLs associated with mean number of leaves **b** warping functions of normalised luminescence of RILs with Tnz-1 alleles and Ws-2 alleles at the SNP1104 locus during a short day to long day photoperiod shift. **c** Boxplots of mean number of leaves for the two main loci for RILs containing the Ws-2 and Tnz-1 copy of the allele. **d** and **f** are equivalent to **a** and **c** but highlighting QTLs associated with the standard error of number of leaves. **e** Normalised luminescence of RILs velocity curves with Tnz-1 alleles and Ws-2 alleles at the SNP42 locus during a short day to long day photoperiod shift.

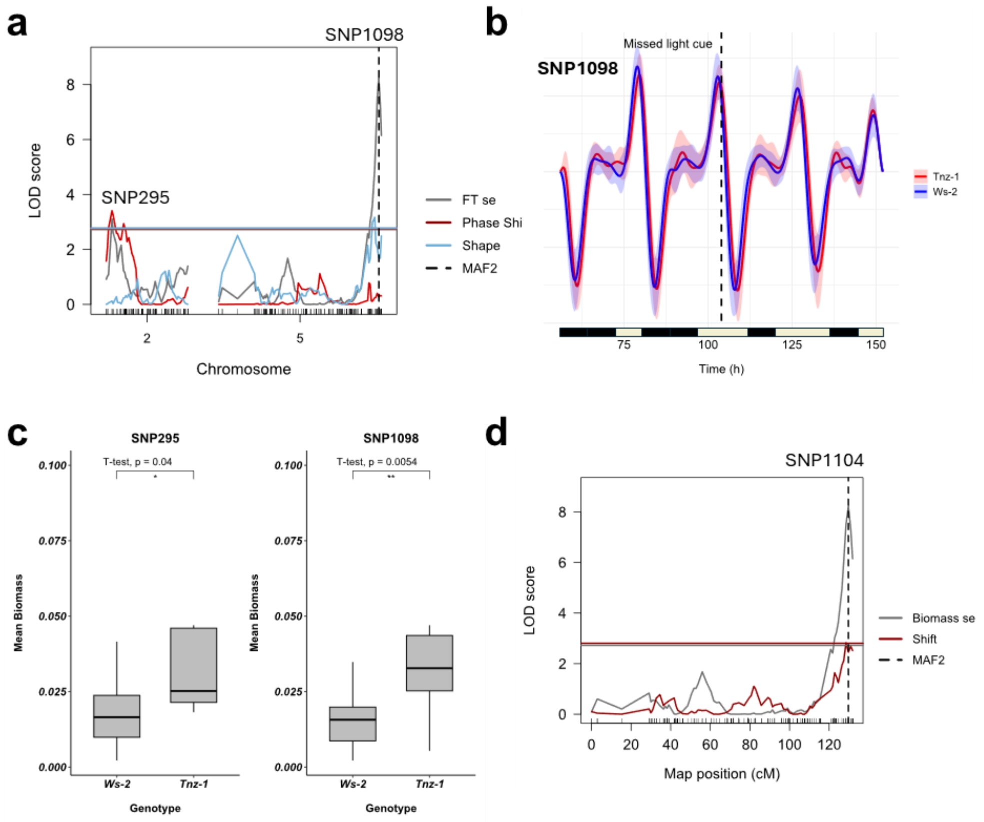


**Supplementary Fig. 22: a** LOD score profile of QTLs associated with mean dry biomass, horizontal line shows 5% threshold. **b** Normalised luminescence of RILs velocity curves with Tnz-1 alleles and Ws-2 alleles at the SNP1098 locus during a short day to long day photoperiod shift. **c** Boxplots of mean dry biomass for the two main loci for RILs containing the Ws-2 and Tnz-1 copy of the allele. **d** is equivalent to **a** but highlighting QTLs associated with the standard error of dry biomass. This is the same region as shown in Fig. 2. See Fig. 2**e** for warping functions of normalised luminescence of RILs with Tnz-1 alleles and Ws-2 alleles at the SNP1104 locus during a short day to long day photoperiod shift. See Figure 2**f** for boxplots of mean dry biomass for the two main loci for RILs containing the Ws-2 and Tnz-1 copy of the allele.

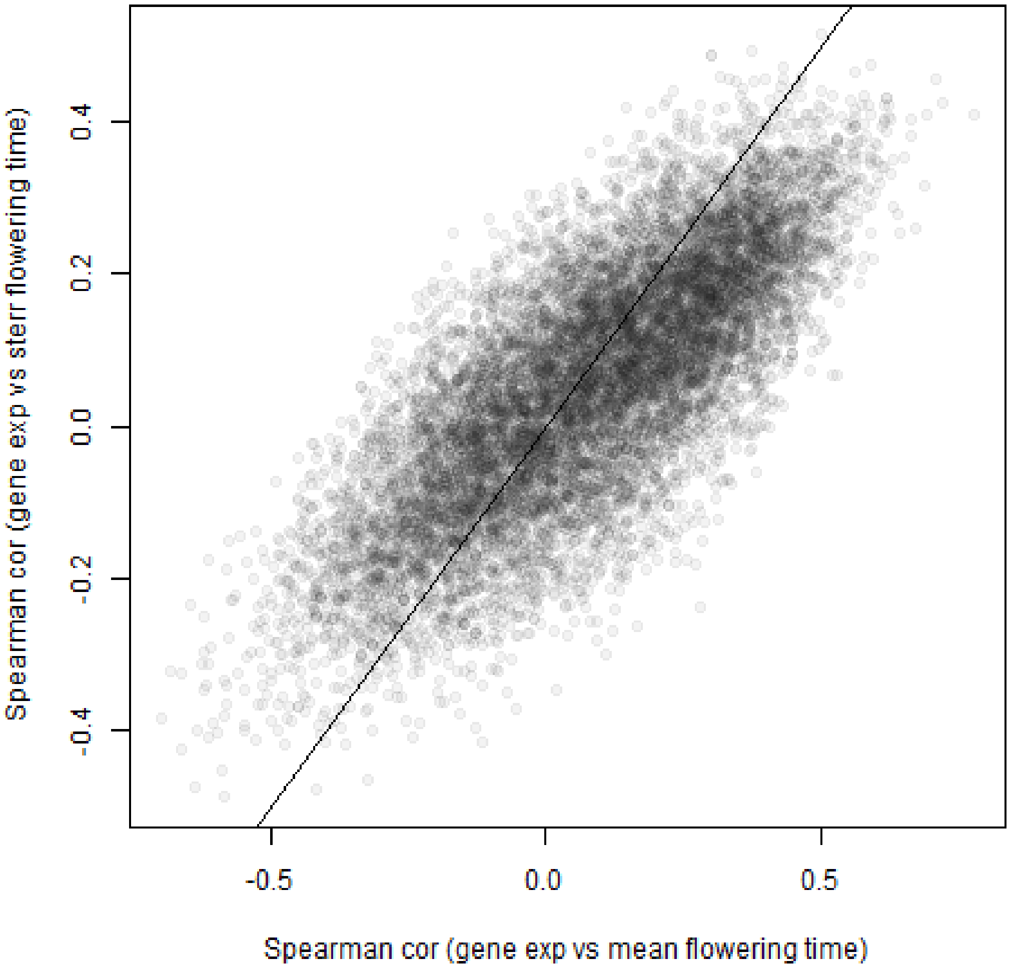


**Supplementary Fig. 23**: For each gene with mean expression >10TPM across all RNA-seq samples, we calculated the Spearman correlation between the gene expression (in TPM) in the control treatment (dark) and either the mean flowering time (x-axis) and the standard error of flowering time (y-axis). A line with x-intercept of 0 and slope of 1 is shown. Even though there is only a weak correlation between mean and standard error of flowering time (Fig S3), we found that most genes whose expression was correlated with mean flowering time were also correlated with standard error of flowering time.
