## Supplementary material for "The genetic basis for synchronised time perception in plant populations": Methods

### Generation of RILs

Transformation of parental accessions Ws-2 and Tnz-1 with CCR2:LUC promoter:reporter construct was the first step in the generation of RIL sets. After initial transformation, a single 'T1' transformant for each accession expressing CCR2:LUC was selected and backcrossed to the respective parent (wild type). This "cleared" the genome for multiple transgene insertions and residual mutations. A single insert line for each parent was then isolated and employed in a full diallel crossing strategy[1,2]. Pair-wise crosses were made among parental accessions, the resultant RIL population obtained Ws-2 X Tnz-1.

The RIL was further comprised of various sub-populations. The sub-populations were categorized based on the crossing approach followed. The crossing methods used were (i) backcross (BC), (ii) F2 [3,4]. The detail procedure adopted to generate the RIL set based on these crossing methods is as follows (**Supplementary Fig. 1**). In BC sub-population, the parent one (Tnz-1, P1-CCR2: pollen donor parent harbouring CCR2:LUC) was crossed to parent two (Ws-2, P2-WT: wild type without CCR2:LUC) to obtain F1 (harbouring CCR2:LUC). Many of these F1 plants were then backcrossed to parent two (Ws-2, P2- WT: wild type without CCR2:LUC) that resulted in a BC1 population. The progeny of BC1 produced BC1F2 plants, from those several lines were selected and self-fertilized for 3-4 generation by SSD to obtain BC1F5 or BC1F6 lines. After each round of selfing, the plants were confirmed for the presence of CCR2:LUC by selection on antibiotics and by imaging for bioluminescence. The resultant BC1F5 or BC1F6 lines were designated as RILs because of their expected genome homozygosity of ~95%. In F2 sub-population, the parent one (Tnz-1, P1-CCR2: pollen donor parent harbouring CCR2:LUC) was crossed to parent two (Ws-2, P2-WT: wild type without CCR2:LUC) to obtain F1 (harbouring CCR2:LUC). Many of these F1 plants were then selfed to obtain F2 segregating progeny. Several F2 lines were selected and self-fertilized for 3-4 generation by SSD with selection for CCR2:LUC. The resultant F5 or F6 lines were designated as RILs (**Supplementary Fig. 1**)

### Agrobacterium mediated transformation of Arabidopsis (Floral Dipping)

All Arabidopsis transformations were performed using a simplified floral-dipping protocol5. Briefly, Agrobacterium strain ABI[6] bearing the required transgene was streaked on YEBS growth media containing appropriate antibiotics. After two days, a starter culture of 25ml YEBS with appropriate antibiotics was inoculated with 2-3 colonies. The starter culture was then incubated at 28°C for two more days, until optimum bacterium growth was achieved. On the day of the transformation, the starting culture was diluted in 500 mL of YEBS and grown for another 6-8 hours. Then, 80 µl of Silwet L-77 was added to the culture. Arabidopsis plants at flowering-initiation stage were submerged for ~20 seconds in the bacterial culture. After dipping, plants were wrapped with plastics bags for 12-18 hours and then transferred to the greenhouse until seeds were matured. T1 transgenic plants were selected MS plates supplemented with appropriate antibiotics.

### Flowering time experiments

Seeds were surfaced sterilised and suspended in 0.1% agar and stratified for 4 days. Seeds were then placed into soil mix and were subjected to 8h photoperiods at 23°C after 7 days, they then had 6 weeks of vernalisation at 4°C and 8-h photoperiods. Post vernalisation to 8h photoperiods at 23°C resumed until all individuals had flowered. Plants were grown in 4 cm-diameter soil-filled pots and bottomed watered two times per week or once per week during vernalisation. The experiment was repeated twice and shown that there was no significant result between experiments, so data was pooled together (**Supplementary Fig. 2**). For up to 40 individuals from each genotype, the number of days post-vernalization to the production of a visible bolt, rosette leaf number at bolting, and dry biomass at bolting were used for this study (**Supplementary Table 1)**.

**Luciferase assays**

Seeds were surface-sterilized and plated onto MS medium with 3% sucrose, and then stratified for 4 days. After stratification, seedlings were entrained under either 8/16 LD or 16/8 LD cycles with a constant temperature of 22° for 7 days. On day 6, seedlings were transferred to black 96-well Microplates with MS medium containing 3% sucrose. Plants were superficially treated with 15 µl 5 mM D-Luciferin. Seedlings were then re-entrained for 1 day under the respective entrainment conditions before being transferred to the TOPCOUNT (Perkin-Elmer [Perkin-Elmer-Cetus], Norwalk, CT). All TOPCOUNT experiments were carried out under blue–red light and a constant temperature of 21°. Experiment were either kept at entrainment conditions throughout or on Day 12 photoperiod was switched from 8/16 LD to 16/8LD, luminescence output was measured until Day 16.

**Functional Data Analysis**

Firstly, the raw topcount files were converted into tables with the time of each measurement and the luminescence readings. Starting with preprocessing of the data the initial aim was to filter curves specifically from ZT56 to ZT152 this provided a 48-hour window of data either side of the perceived shift. Curves also had to exceed a minimum luminescence threshold set to remove plants that had died (100). It was also important to identify and remove plants that would initially have been luminescing but then died making them unsuitable for further analysis. To achieve this, we focused on the 48 hours post perceived shift detrended and a Lomb-Scargle periodogram analysis is performed to identify peaks indicative of periodicity. Curves failing to meet the significance threshold (0.001) in the periodogram analysis are classed at non-rhythmic removed from the dataset.

The next step was to estimate the discrete curves into smooth continuous curves using Functional Data Analysis (FDA) methods[7–10]. Initially, luminescent curve was normalised to the range [0, 1]. Then each individual curve using its corresponding specific time values using basis functions were defined via B-splines and setting a penalty for roughness and functional parameters are generated. Here the number of basis functions were maximised as there was no worry in overfitting as each curve was being estimated individually, lambda was used to smooth the functions. Outliers in this data frame were detected using FDA outlier detection methods (‘fdaoutlier’), specifically the Total Variation (TV) Smoothing and Derivative-based Measures of Scale (MSS)[11]. Detected outliers were then removed from further analysis. Next, a new time vector was created to cover the range of all original time values. Each curve was evaluated under this new time vector, resulting in a matrix of all curves under the same time points.

Further analysis was conducted whereby genotype specific functional traits were identified. Firstly, functional principal component analysis (FPCA) was completed for the curves in all conditions in both the original curves and the first and second derivatives. See **Supplementary Fig. 15** as example of first derivative. The resulting FPCA's there are different categories of traits Shape, Spread and Shift related. Here the mean position of each genotype was determined (Shape) also the Euclidean distances for each individual to conform to the mean (Spread of Shape, **Supplementary Fig. 16**). Additionally, each individual curves could be estimated for pre and post perceived shift (in the SDSD and LDLD condition there was no shift, **Supplementary Fig. 17**). The distance each individual and genotypes moved within the PC1, PC2, and PC3 space was recorded and used as a QTL trait. Lastly using curves as functions each curve was able to be mapped to the mean curve for each genotype. This was evaluated pre and post shift using time warping function, this allowed for the assessment of the change in shape of curves as well as the variation present either side of the shift. For full information of functional traits see **Supplementary Table 5**.

**RNA-seq experiment**

Seeds were surface-sterilized and plated onto MS medium with 3% sucrose in clusters of ~50 seeds and then stratified for 4 days. After stratification, seedlings were entrained in two cabinets under 8/16 LD cycle with a constant temperature of 22° for 12 days. On day 12, individuals in treatment cabinet were shifted to 16/8 LD. On day 12 at ZT10 all plants were harvested and snap-frozen in liquid nitrogen.

**RNA-seq data analysis**

Total RNA was isolated from whole plant tissue using the Qiagen RNeasy Plant Mini Kit. Residual genomic DNA was removed using the Invitrogen Turbo DNA-free kit, according to the manufacturer's protocol. Libraries were prepared with the NEBNext Ultra II Directional Library Prep Kit for Illumina, using the NEBNext poly(A) magnetic isolation module. Quality control was performed with the Agilent 2100 Bioanalyzer instrument (Part no. G2939BA). Finally, a total of 130 libraries were sequenced, via Novagene, using one lane on an Illumina NovaSeq system.

Before analysis of the raw sequencing data, FastQC v0.11.7 was used to assess read quality[12]. Illumina adapters were trimmed using CutAdapt v3.4[13]. Reads were quantified using Salmon v1.6.0 and the TAIR10 transcriptome [14,15]. Variant calling was done using GATK workflow described on the GATK website (https://gatk.broadinstitute.org).

For further analysis, transcripts per million (TPM) was used as the measure of relative gene expression across samples (**Supplementary Table 2**). TPM values were filtered to remove genes with very low expression, to avoid biassing clustering and other analyses. Genes were removed for low expression if they TPM below 10 in 50% of samples. This left 11762 genes for further analysis.

**Genetic map construction**

The genetic map for Ws-2 × Tnz-1 recombinant inbred line (RIL) population, built on RNA-seq data was constructed using the following methods. A combined VCF files was generated containing each sample in the control condition. Since not all SNPs are found in all genotypes, this was filtered for SNPs present in >50% of individuals. From this unique list, information regarding the position in base pairs and the chromosome location of each SNP was retrieved and further filtered for only biallelic SNPs differing across parental lines and exceeding a quality score of 30.

In order to get a more reliable genotypic score, cancelling out any SNPs miscalls, and to reduce the overall number of markers, SNPs were grouped into bins. 1125 artificial bins of 100 kbp were created along the whole genome, in a few cases where there were no SNPs passing all criteria these bins could be larger. The scoring of the genotype was obtained based on the SNP information within each bin. For regions at the transition between two genotypic blocks, the bin score was rounded up and assigned to the closest genotypic score.

The bins are ordered based on the genome sequence, each bin is used as a marker and the midpoint position of the bin is used as the marker position. Markers were named SNP for SNP marker, followed by the number of the bin. As an example SNP1 corresponds to the first SNP marker at 0.05 Mbp on chromosome 1. Based on a 100-kbp window SNP binning method, 789 bin-markers were identified, physically anchored on the genome. The total length of the RNA-seq genetic map spans 471.80 centimorgans (cM) with an average marker distance of 0.6 cM and a maximum marker distance of 13.80 cM. The genetic distances were estimated using the "est.map" function with "kosambi" distance from the R/qtl package[16,17]. The correct order of the markers was verified by pairwise marker linkage analysis using the "est.rf " function. The recombination rate was determined based on the linear relation between the genetic and the physical positions of the marker. The segregation pattern was tested for all markers to identify markers that show significant distortion at the 5% level, after a Bonferroni correction for multiple testing. The genetic map and genotypic data are available in **Supplementary Table 3**.

**QTL and eQTL analysis**

Traits for QTL analysis were derived from the flowering time experiment (developmental) and the FDA output (functional). QTL mapping was conducted using the R/qtl (v) package16. Genotype and phenotype data were integrated, and a genome scan was performed using the Haley-Knott (HK) regression method[18,19]. To determine the significance thresholds for QTL detection, 1,000 permutations of the data were performed for each trait, and genome-wide significance thresholds were set at the 95th percentile of the permutation distribution. QTLs that exceeded the permutation threshold were further analysed to identify overlaps among traits. Only QTLs shared between developmental traits and other traits of interest were retained for subsequent analysis (**Supplementary Table 5**).

**Elastic net**

TPM values were z-scored before input. A set of genes was curated by selecting genes differentially expressed across conditions between parents, these were used as the set of possible predictors. A tiered cross-validation approach was used to find optimal hyperparameters. This was implemented using `scikit-learn` in Python v3.10.4 using the `ElasticNetCV` function[21]. Firstly, leave-one-out-cross-validation (LOOCV) was used to produce a 63 separate test-train splits, where the testing set only consisted of a single sample. Then for each training set, 5-fold cross validation was used to pick optimal `alpha` and `l1_ratio`, with possible inputs for `l1_ratio` of 0.01, 0.5, 0.7, 0.9, 0.95, 0.99, and 1.0. The output of this process was a set of optimal hyperparameters and a trained Elastic Net model for each test-train splits. The exact coefficients in the models are summarised in **Supplementary Table 4**.

**Analysis of splice junctions**

Splice junctions were extracted from the STAR alignment SJ.out file. Initially, all splice junctions that were found within the genome coordinates of each of the FLC family genes (AT5G10140, AT1G77080, AT5G65050, AT5G65060, AT5G65070, AT5G65080) were extracted (not including the trans-gene splice junctions). Splice junctions that were supported by more than 2 reads were identified. Of those, only splice junctions that constituted at least 0.1% of the remaining splice sites were considered. Clustering was performed using hierarchical clustering, using the default parameters in the pheatmap function in R[22]. Two main clusters of splice sites were identified. For visualisation purposes, we wished to display fewer rows in **Fig. 3a**, so we performed a t-test between the two clusters for each of the splice junction read counts and chose to display all rows that had unadjusted p-values <0.05. Coverage over exons was visualised using the Integrated Genome Viewer (IGV)[23]. Trans-splice sites were then extracted from the SJ.out file from the STAR aligner (**Supplementary Table 6**).

**Analysis of 1001 genome project data**

The pseudogenome programme in the 1001 genome project website was used (https://tools.1001genomes.org/pseudogenomes/, November 2024[24]) to extract the sequences of KH17 (AT3G32940) and KH29 (AT5G56140) in fasta format. These sequences were aligned using Clustal Omega (https://www.ebi.ac.uk/jdispatcher/msa/clustalo, November 2024[25]), and were loaded in R using the seqinr package. The conserved domains were identified using blastx (https://blast.ncbi.nlm.nih.gov/, November 2024[26]) and the structures were visualised with Alphafold (https://alphafold.com/, November 2023[27]). To calculate the rolling average % identity, the total number of sequences that matched the consensus sequence (i.e. the most common base) were calculated and divided by the total number of sequences (including gaps and/or Ns), and then this was averaged over a 10 base-pair window. To calculate the mutational hotspots, the % identity was calculated by finding matches to the consensus and dividing by the total number of sequences, but this time *without* including gaps and/or Ns. Any base with less than 95% identity was identified as a mutational hotspot. Variant clustering was performed with pheatmap and depicted in a map using the maps package[22]. Flowering time experiments were performed using the same protocol as the RILs and in the same growth chambers (**Supplementary Table 7**).

**Methods References**


1. Blanc, G., Charcosset, A., Veyrieras, J.-B., Gallais, A. & Moreau, L. Marker-assisted selection efficiency in multiple connected populations: a simulation study based on the results of a QTL detection experiment in maize. *Euphytica* **161**, 71–84 (2008).
2. Rebai, A. & Goffinet, B. Power of tests for QTL detection using replicated progenies derived from a diallel cross. *Züchter Genet. Breed. Res.* **86**, 1014–1022 (1993).
3. Kover, P. X. *et al.* A Multiparent Advanced Generation Inter-Cross to fine-map quantitative traits in Arabidopsis thaliana. *PLoS Genet.* **5**, e1000551 (2009).

4. Lee, M. *et al.* Expanding the genetic map of maize with the intermated B73 × Mo17 (IBM) population. *Plant Mol. Biol.* **48**, 453–461 (2002).
5. Davis, A. M., Hall, A., Millar, A. J., Darrah, C. & Davis, S. J. Protocol: Streamlined sub-protocols for floral-dip transformation and selection of transformants in Arabidopsis thaliana. *Plant Methods* **5**, 3 (2009).
6. Schomburg, F. M., Patton, D. A., Meinke, D. W. & Amasino, R. M. FPA, a gene involved in floral induction in Arabidopsis, encodes a protein containing RNA-recognition motifs. *Plant Cell* **13**, 1427–1436 (2001).
7. Ramsay, J. O., Wickham, H., Ramsay, M. J. O. & deSolve, S. Package "fda." (2024).
8. Wang, J.-L., Chiou, J.-M. & Müller, H.-G. Functional data analysis. *Annu. Rev. Stat. Appl.* **3**, 257–295 (2016).
9. Levitin, D. J., Nuzzo, R. L., Vines, B. W. & Ramsay, J. O. Introduction to functional data analysis. *Can. Psychol.* **48**, 135–155 (2007).
10. Ramsay, J. O., Hooker, G. & Graves, S. Introduction to functional data analysis. in *Functional Data Analysis with R and MATLAB* 1–19 (Springer New York, New York, NY, 2009).
11. Febrero–Bande, M. & Fuente, M. D. L. Statistical computing in functional data analysis: The R package fda.Usc. *Journal of Statistical Software* **051**, 1–28 (2012).
12. Wingett, S. W. & Andrews, S. FastQ Screen: A tool for multi-genome mapping and quality control. *F1000Res.* **7**, 1338 (2018).
13. Martin, M. Cutadapt removes adapter sequences from high-throughput sequencing reads. *EMBnet.journal* **17**, 10–12 (2011).
14. Patro, R., Duggal, G., Love, M. I., Irizarry, R. A. & Kingsford, C. Salmon provides fast and bias-aware quantification of transcript expression. *Nat. Methods* **14**, 417–419 (2017).
15. Berardini, T. Z. *et al.* The Arabidopsis information resource: Making and mining the "gold standard" annotated reference plant genome: Tair: Making and Mining the "Gold Standard" Plant Genome. *Genesis* **53**, 474–485 (2015).

16. Broman, K. W., Wu, H., Sen, S. & Churchill, G. A. R/qtl: QTL mapping in experimental crosses. *Bioinformatics* **19**, 889–890 (2003).
17. Arends, D., Prins, P., Jansen, R. C. & Broman, K. W. R/qtl: high-throughput multiple QTL mapping. *Bioinformatics* **26**, 2990–2992 (2010).
18. Knott, S. & Haley, C. Aspects of maximum likelihood methods for the mapping of quantitative trait loci in line crosses. *Genetics Research* **60**, 139–151 (1992).
19. Feenstra, B., Skovgaard, I. & Broman, K. Mapping quantitative trait loci by an extension of the Haley–Knott regression method using estimating equations. *Genetics* **173**, 2269–2282 (2006).
20. Shabalin, A. Matrix eQTL: ultra fast eQTL analysis via large matrix operations. *Bioinformatics* **28**, 1353–1358 (2011).
21. Pedregosa, F. *et al.* Scikit-learn: Machine learning in Python. *the Journal of machine Learning research* **12**, 2825–2830 (2011).
22. Kolde, R. & Kolde, M. R. Package "pheatmap." *R package* (2015).
23. Thorvaldsdóttir, H., Robinson, J. T. & Mesirov, J. P. Integrative Genomics Viewer (IGV): high-performance genomics data visualization and exploration. *Brief. Bioinform.* **14**, 178–192 (2013).
24. Weigel, D. & Mott, R. The 1001 genomes project for Arabidopsis thaliana. *Genome Biol.* **10**, 107 (2009).
25. Sievers, F. & Higgins, D. G. Clustal omega. *Curr. Protoc. Bioinformatics* **48**, 3.13.1-3.13.16 (2014).
26. James Kent, W. BLAT—The BLAST-Like Alignment Tool. *Genome Res.* **12**, 656–664 (2002).
27. Jumper, J. *et al.* AlphaFold 2. Preprint at https://predictioncenter.org/casp14/doc/presentations/2020_12_01_TS_predictor_AlphaFold2.pdf.
